# Properties of genes encoding transfer RNAs as integration sites for genomic islands and prophages in *Klebsiella pneumoniae*

**DOI:** 10.1101/2020.11.02.365908

**Authors:** Camilo Berríos-Pastén, Rodolfo Acevedo, Patricio Arros, Macarena A. Varas, Kelly L. Wyres, Margaret M. C. Lam, Kathryn E. Holt, Rosalba Lagos, Andrés E. Marcoleta

## Abstract

The evolution of traits including antibiotic resistance, virulence, and increased fitness in *Klebsiella pneumoniae* and related species has been linked to the acquisition of mobile genetic elements through horizontal transfer. Among them, genomic islands (GIs) preferentially integrating at genes encoding tRNAs and the tmRNA (t(m)DNAs) would be significant in promoting chromosomal diversity. Here, we studied the whole set of t(m)DNAs present in 66 *Klebsiella* chromosomes, investigating their usage as integration sites and the properties of the integrated GIs. A total of 5,624 t(m)DNAs were classified based on their sequence conservation, genomic context, and prevalence. 161 different GIs and prophages were found at these sites, hosting 3,540 gene families including various related to virulence and drug resistance. Phylogenetic analyses supported the acquisition of several of these elements through horizontal gene transfer, likely mediated by a highly diverse set of encoded integrases targeting specific t(m)DNAs and sublocations inside them. Only a subset of the t(m)DNAs had integrated GIs and even identical tDNA copies showed dissimilar usage frequencies, suggesting that the genomic context would influence the integration site selection. This usage bias, likely towards avoiding disruption of polycistronic transcriptional units, would be conserved across Gammaproteobacteria. The systematic comparison of the t(m)DNAs across different strains allowed us to discover an unprecedented number of *K. pneumoniae* GIs and prophages and to raise important questions and clues regarding the fundamental properties of t(m)DNAs as targets for the integration of mobile genetic elements and drivers of bacterial genome evolution and pathogen emergence.

## BACKGROUND

The Gram-negative bacterium *Klebsiella pneumoniae* has been traditionally considered as an opportunistic pathogen causative of nosocomial infections [1]. However, an increasing number of severe community-acquired infections caused by hypervirulent *K. pneumoniae* strains are being reported globally [2,3]. On the other hand, a distinct set of lineages of *K. pneumoniae* strains resistant to multiple antimicrobials have emerged and caused several outbreaks in diverse geographical locations. Furthermore, this species is considered a major source and a key trafficker of antibiotic resistance genes that can spread among other Gram-negative pathogens [4,5]. This has led to multidrug-resistant *K. pneumoniae* being declared as an urgent threat by the U.S. Centers for Disease Control and Prevention (CDC) and the World Health Organization (WHO). Genomic analyses of hypervirulent and multi-resistant strains revealed that these traits are mainly determined by specific sets of genes acquired horizontally, which are encoded in a range of mobile elements that would mediate their dissemination [6,7]. In particular, it has been reported that several genomic islands of this species carry a wide repertoire of virulence and drug-resistance associated factors [8–10].

Genomic islands (GIs) are discrete DNA segments variably present in equivalent positions of the chromosome. In certain conditions, some of these excise forming a circular intermediate that could be transferred to a different host and integrate into a new site, mainly at genes encoding transfer RNAs (tDNAs) or the transfer-messenger RNA (tmDNA, also named *ssrA*) [11–13]. For convenience, “t(m)DNA” will be used when speaking of both kinds of genes. GIs can carry hundreds of cargo genes encoding a variety of functions and accessory metabolic pathways. Some of them, frequently referred to as integrative conjugative elements (ICEs), encode a type-IV secretion system and include a transfer origin (oriT) sequence allowing the spread of the element through conjugation [14]. Additionally, GIs often comprise a gene coding for an integrase that mediates the integration and excision of the element, normally recognizing the 3’ end of the target t(m)DNA (sequence known as attB) and catalyzing the site-specific recombination with an identical sequence present in the GI (attP). Upon integration, the attB site is duplicated resulting in a direct repeat that delimitates both ends of the integrated GI [11]. It is currently accepted that the choice of the integration site largely depends on the integrase protein, which has evolved to recognize a specific attB sequence and thus a defined t(m)DNA. Although the usage of t(m)DNAs as integration sites for GIs and prophages has been documented for different bacterial species, the reason that these genes are preferred over other genes is still an open question. Moreover, to our knowledge, no previous studies have investigated the properties of the complete set of t(m)DNAs typically present in the chromosome of a given bacterial species, as integration sites for these elements.

In a previous study, we analysed 50 *K. pneumoniae* chromosomes, leading to the identification of 12 GI families that integrate into asparagine tDNAs, proposing these genes as hotspots for GI integration in this species [8]. Most of the identified islands encoded factors with proven or potential roles in pathogenesis and drug resistance. Among them, we found the previously described ICE*Kp1* element [15], harbouring genes for the production of yersiniabactin and salmochelin siderophores (*ybt* and *iro* genes, respectively), the *rmpA* regulator of capsule and hypermucoviscosity, as well as genes for conjugal transfer. Additionally, we characterized the GIE492 island, encoding the determinants for the production of the antimicrobial peptide microcin E492 and salmochelin (*mce* genes). Remarkably, we found that GIE492 and other GIs carry a cryptic oriT highly identical to that present in ICE*Kp1*, suggesting that these are mobilizable GIs that could take advantage of the conjugative machinery encoded by ICE*Kp1* and related elements including the additional 13 known *ybt* carrying ICE*Kp*s (designated ICE*Kp2*-ICE*Kp14*) [9]. We showed that GIE492 was highly prevalent among *K. pneumoniae* strains isolated from liver abscesses. This observation was further corroborated upon the detailed genomic analysis of the clonal group 23 (by far the most common clonal group identified among hypervirulent *K. pneumoniae* infections globally), which indicated that more than 90% of the isolates from this group harbour the GIE492, along with an ICE*Kp* element (mainly ICE*Kp10*), and each of these integrated into an asparagine tDNA [16]. All this evidence supports the major role of GIs in the evolution and pathogenesis of *K. pneumoniae*. Furthermore, we observed that a typical *K. pneumoniae* chromosome harbours four copies of the asparagine tDNA, each of them located in a specific and highly conserved genomic context. Although the copies were 100% identical (and thus offer the same attB site), they were occupied by GIs with a very dissimilar frequency, suggesting that unattended factors could influence the selection of the integration site [8]. This observation was further corroborated after studying the presence of different ICE*Kp* variants among a set of over 2,000 *K. pneumoniae* isolates, where the integration frequency of these elements in each of the four asparagine tDNAs was very dissimilar [9]. In this regard, it remains unknown if the usage bias among identical copies of asparagine tDNAs extends to other tDNAs, as well as the possible factors that account for this phenomenon. It is also unknown if all or only a subset of the t(m)DNAs from the *K. pneumoniae* chromosome are used as GI integration sites. Additionally, the analysis of the whole set of KpSC t(m)DNAs as integration sites should lead to the discovery of a significant number of new GIs, allowing further investigation of their potential role in clinically relevant traits.

In this study, we investigated the prevalence, sequence conservation, genomic context, and usage as integration sites of the t(m)DNAs from a set of 66 chromosomes representative of diverse lineages from *K. pneumoniae* and closely related species. Also, we studied the features and encoded functions of the GIs and prophages integrated at these sites, the role of the integrases and other possible factors in integration site selection and usage frequency, and possible evidence of GI and prophage dissemination through horizontal gene transfer.

## RESULTS

### Identification, classification, and nomenclature of *K. pneumoniae* t(m)DNAs

We studied the t(m)DNAs present in the chromosomes of 66 strains from four species of the *K. pneumoniae* species complex (KpSC), namely, *Klebsiella pneumoniae sensu stricto* (previously known as phylogroup Kp1), *Klebsiella quasipneumoniae* subsp. *quasipneumoniae* (Kp2), *Klebsiella quasipneumoniae* subsp. *similipneumoniae* (Kp4), and *Klebsiella variicola* subsp. *variicola* (Kp3) [17,18]. These strains are of diverse multi-locus sequence types [19], were isolated from a variety of niches, collected at diverse times, and have distinct virulence and drug resistance potential (Supplementary Table 1). Using the tool ARAGORN [20], followed by intensive manual curation, we identified a total of 5,624 tDNAs among our dataset, ranging from 77 to 92 tDNAs per chromosome (mode=86 tDNAs) (Figure 1A). The length of the tDNAs varied between 73 bp to 94 bp (mode=75 bp) (Figure 1B). All chromosomes had a single copy of *ssrA* (length=363 bp).

**Figure 1.**
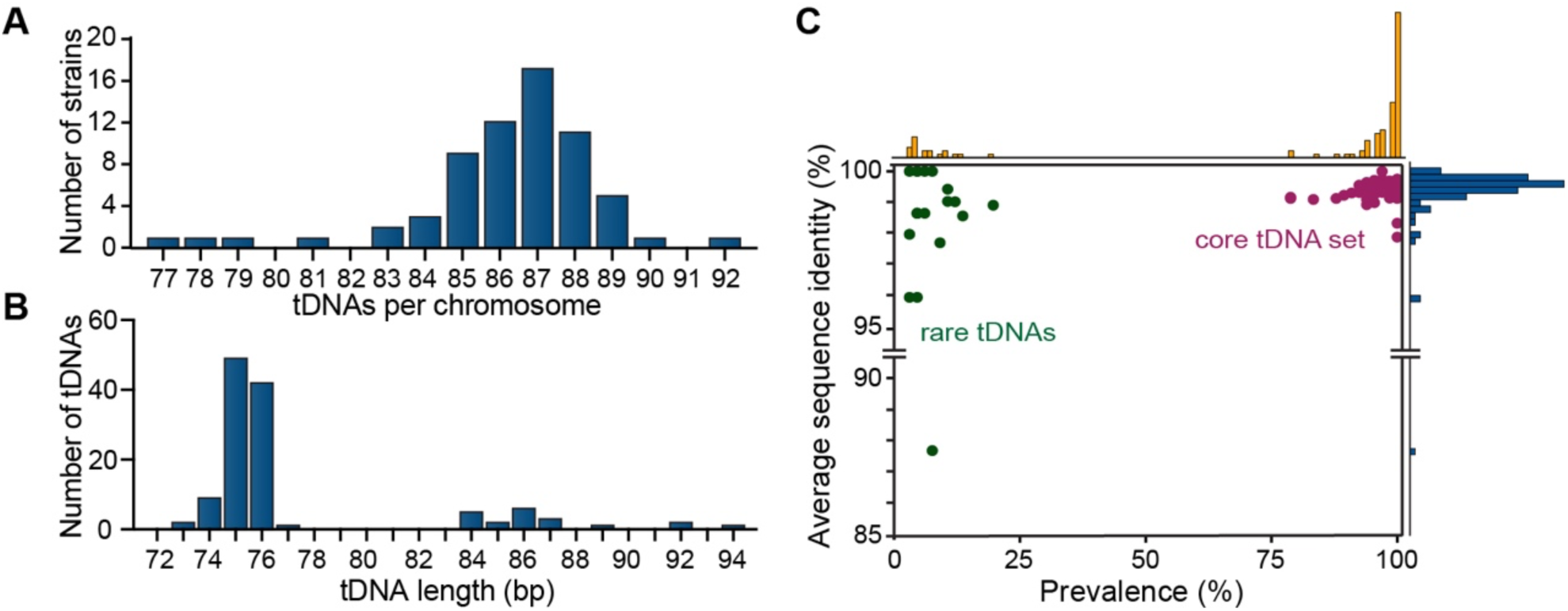
General features of the KpSC tDNAs. tDNAs from a set of 66 KpSC strains were identified and classified. (A) Frequency distribution of the total number of tDNAs per strain. (B) Length frequency distribution of the identified tDNAs. (C) Average sequence identity shared by all the tDNAs carrying the same anticodon(vertical axis), and their prevalence (horizontal axis), calculated as the percent of strains harbouring each Tdna class. Bar charts indicate frequencies of t(m)DNA genes.

Next, we aimed to classify all t(m)DNAs and compare them among the different KpSC chromosomes. However, the original annotation of these genes varied significantly across the different genomes. Moreover, the bioinformatics tools commonly used for tDNA annotation in bacterial genomes, including ARAGORN and tRNAscan-SE [21], name each tDNA only specifying the amino acid charged by the encoded tRNA and the anticodon sequence (e.g. asn-GTT). This partial and variable tDNAs annotation is mainly due to the lack of consensus regarding how to annotate these genes in bacteria, and as a major drawback, it does not allow differentiation of identical tDNA copies located in different positions of the chromosome (different genomic context). Given that the genomic context can influence the usage of tDNAs as GI integration sites, we developed a new nomenclature system for naming these genes in strains from KpSC, which can be adapted to other bacterial species. The system assigns to each tDNA a name composed of (i) the three-letter code of the amino acid transferred by the tRNA (in lowercase) followed by (ii) an Arabic numeral that is different for each anticodon specifying the same amino acid; and (iii) a capital letter that conveys the specific genomic context where the tDNA can be found (for more details see Supplementary Figure 1). To facilitate tDNA annotation following this new nomenclature in KpSC chromosomes, we developed a tool named Kintun-VLI (available at https://github.com/GMI-Lab/Kintun-VLI). The name *ssrA* was retained for the tmDNA [13]. Using this new nomenclature system, we reclassified our set of 5,624 tDNAs into 115 non-redundant genes based on their anticodon and characteristic genomic context. This classification scheme allowed us to study several properties of the KpSC t(m)DNAs, as described in the following sections.

### Prevalence and nucleotide conservation of the KpSC t(m)DNAs: defining the core set

Following t(m)DNA classification, we determined their prevalence and nucleotide sequence conservation. 86 of the 115 different tDNAs were found in more than 75% of the strains, constituting the core tDNA set (Figure 1C). The remaining tDNAs had a prevalence <25% and were mainly located inside putative mobile elements or regions showing uncommon gene organization, which in some cases may be the result of assembly errors. Sequence conservation of each different tDNA across all strains was evaluated through multiple alignments using ClustalW [22]. We found a mean identity *≥*98% for each tDNA from the core set (Figure 1C), while lower identity values were observed for some of the rare tDNAs. As mentioned, a single copy of *ssrA* was found in all strains, showing a mean identity of 99.9%. Therefore, only tDNAs belonging to the core set and *ssrA* were included in subsequent analyses. These include 40 tDNAs encoding distinct anticodons, 19 of them present in 2 to 7 copies (Supplementary Table 2). As observed for other bacterial species, this set is incomplete i.e. it does not include anticodons matching the 62 possible codons according to the classic genetic code (excluding the stop codons). In this regard, it was proposed that these orphan codons can be translated by some of the encoded tRNAs carrying partially complementary anticodons, as a consequence of post-transcriptional modification and the wobbling effect [23]. The tDNA copy number and distribution observed for the KpSC core set closely resembled that determined for *E. coli* K12 and DSM30080, which in this and other bacteria was demonstrated to have evolved to match the codon usage patterns and increase translation efficiency [24–27]. Indeed, we observed similar codon usage in *K. pneumoniae* compared to *E. coli* (Supplementary Table 2).

### Genomic context of t(m)DNAs and identification of integrated mobile genetic elements

We found that the gene organization of the regions located within 4 kbp upstream of each t(m)DNA was highly conserved across the different chromosomes, sharing >90% average sequence identity in most cases (Supplementary Figure 2). Although to a lesser extent, the downstream regions were also conserved, unless a mobile genetic element had integrated at these sites causing the shift of this region to a position immediately downstream of the element. Through comparing the genomic contexts of each t(m)DNA across our set of chromosomes, we deduced their typical organization in absence of any integrated mobile genetic element, which we refer to as the “virgin t(m)DNA locus”. The different t(m)DNAs were found either adjacent to protein-coding genes, genes encoding rRNAs, or other tDNAs forming operon-like gene clusters (Supplementary Figure 3). From knowing the virgin configuration of each t(m)DNA, we devised a strategy to identify all the putative GIs integrated into these genes based on comparing their contexts with the respective virgin configuration and detection of unshared regions by using BLASTn [28] and ProgressiveMauve [29], followed by intensive manual curation. For these analyses, a GI was defined as a DNA segment variably present adjacent to a t(m)DNAs across different strains. A minimum length of 3.5 kbp was defined to filter out insertion sequences or other forms of small insertions. This threshold also corresponded to the smallest putative GI found harbouring an integrase and direct repeats, which are characteristic features of GIs. We identified 412 putative t(m)DNA-associated GIs among the 66 chromosomes. All strains had four to ten GIs, with a mean of six GIs per chromosome. The total chromosome size observed for all the strains ranged from 5.1 to 5.7 Mbp, where the identified GIs accounted for 1% to 6% of the total length (50-350 kbp). Significant positive correlations were observed between the chromosome length and the number of GIs per chromosome (Figure 2A), the total length corresponding to GIs (Figure 2B), and the total number of GI-encoded genes (Figure 2C), supporting that larger chromosomes harbour a higher GI content. The described approach to identify GIs was further leveraged to develop an algorithm implemented as part of the Kintun-VLI tool, which allows detection of t(m)DNA-associated GIs in a given *K. pneumoniae* chromosome without dependence on the identification of features considered common but not universally present in these elements (e.g. repeats, integrase-coding genes).

**Figure 2.**
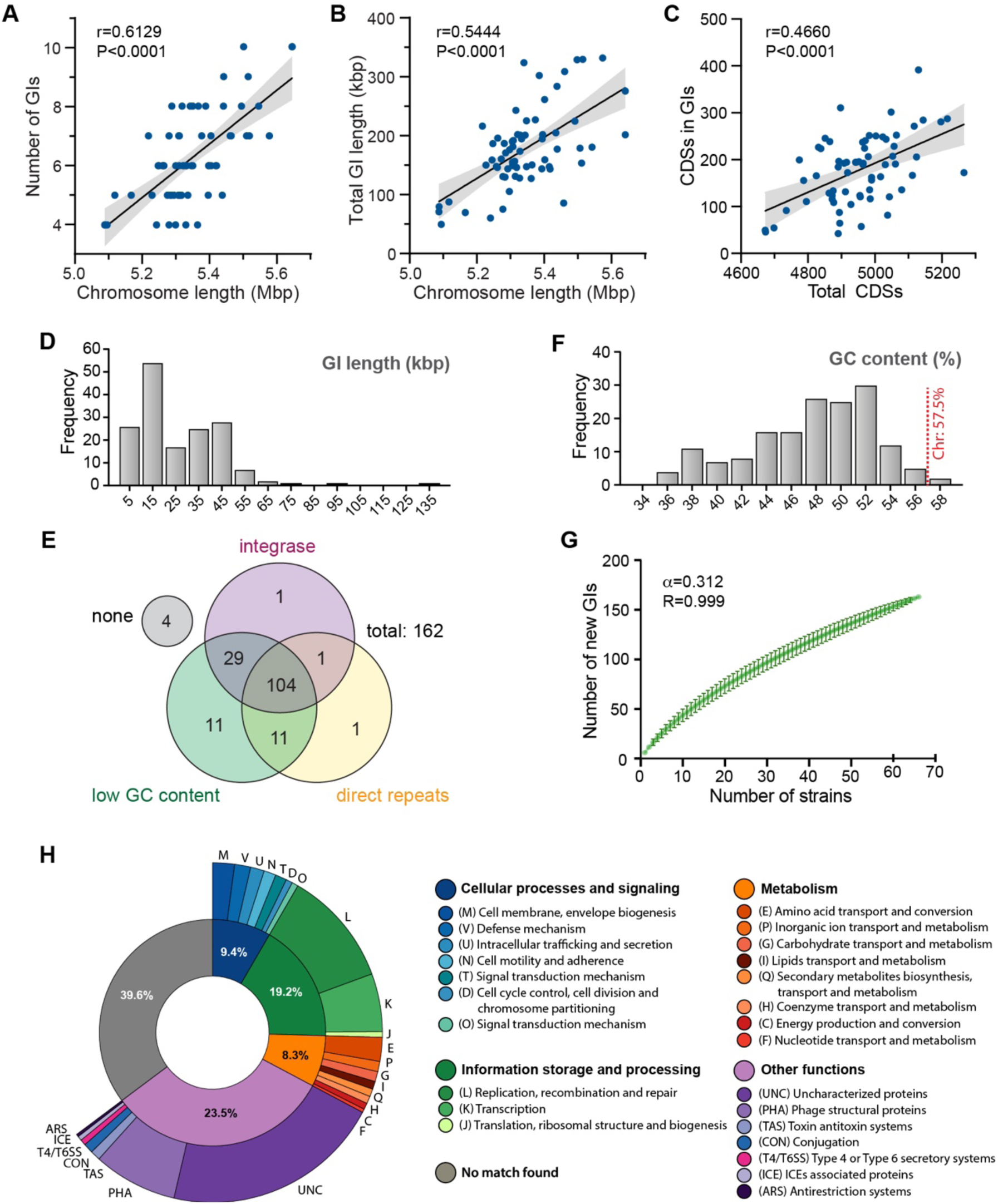
Properties and encoded functions of the KpSC genomic islands. Number of identified t(m)DNA-associated GIs per strain (A) and total length of the chromosome corresponding to GIs (B), as a function of the total chromosome length. The Spearman’s r coefficient and the associated P-values are indicated. Areas shaded in grey correspond to the 95% confidence interval of the linear fit slope. (C) frequency distribution of GI length. (D) Venn diagram showing the number of GIs harbouring an integrase-coding gene, direct repeats, and/or a low GC-content. (E) frequency distribution of the GC content among the detected GIs. (F) amino acid length-frequency distribution of the integrase proteins encoded in the GIs. (G) accumulation curve describing the tendency to discover new GIs while exploring additional *K. pneumoniae* strains. Error bars represent the standard deviation from permutating all the possible orders of strain additions. (H) eggNOG functional categorization of the 3,540 non-redundant gene families encoded in the detected GIs.

### The KpSC chromosomes host a remarkable diversity of t(m)DNA-associated genomic islands and prophages

To further characterize the identified GIs, their sequences were extracted from the chromosomes, aligned, and classified into 162 non-redundant GIs according to sequence similarity (coverage >60%, identity >80%) (Supplementary Table 3). For convenience, each GI was named according to the tDNA where it was found integrated, followed by a Roman numeral distinguishing different GIs that integrate into the same tDNA, as described previously [8]. The size of GIs ranged from *∼*3.5 kbp to *∼*133 kbp (mode=15 kbp) (Figure 2C). Using BLAST-based sequence alignments and custom Python scripts, we searched our non-redundant GI collection for features commonly associated with these elements, namely, flanking direct repeats, low GC content, and the presence of integrase-coding genes. One hundred and four of the 162 GIs showed all these three features, while 4 GIs showed none of them (Figure 2D). A “low GC content” was defined as a value lower than the mean calculated from all the chromosomes (57.50±0.19%) minus five times the standard deviation (<56.55%). Only 7 of the 162 GIs did not meet this criterion, with most of them showing a GC content <52%, going down to *∼*36% (Figure 2E). One hundred and thirty-five GIs harboured at least one gene encoding an integrase protein (Figure 2D) of variable length (mode=420 aa) (Figure 2F and Supplementary Table 3). Additionally, in 117 GIs we identified flanking direct repeats (Figure 2D), most of them including part of the respective t(m)DNAs, with lengths ranging from 15 bp to 221 bp (mode=20-25 bp). Using the tool PHASTER [30], 32 putative GIs were predicted as intact prophages, 8 as questionable prophages, and 10 as incomplete prophages (Supplementary Table 3). Prophages and GIs showed essentially the same structure and features, including comparable direct repeats and integrase proteins, but the former carrying genes related to phage assembly and replication.

To explore the diversity of t(m)DNA-associated GIs and prophages within our sample set and the expected diversity of these elements yet to be discovered, we constructed an accumulation curve (Figure 2G). The factor α, calculated from the adjustment of the curve according to the Heaps’ law where α<1 reflects an open set [31], indicated that analysis of additional strains would lead to the discovery of a significant number of new GIs/prophages (α = 0.312 ± 0.003; R = 0.999). Gene content analysis using Roary [32] indicated a total of 16,161 gene families among the 66 chromosomes, from which 3,564 were shared by at least 95% of the strains (core genome: 22.1%; identity cut-off=95% at protein level). From the 12,597 gene families constituting the accessory genome of this set, 3,540 (28.1%) were encoded inside t(m)DNA-associated GIs, supporting these elements as main actors in mediating chromosomal diversity in KpSC. Remarkably, gene presence/absence analysis indicated that the overall gene content was highly different among the distinct GIs (Supplementary Figure 4). In agreement, a low shared sequence identity was observed after performing pairwise GI alignments using BLASTn. From a total of 13,122 comparisons, only 14 matches (>60% identity; e-value <0.001) were observed between different GIs integrated at the same t(m)DNA, and 26 between GIs integrated at distinct t(m)DNAs (Supplementary Figure 5). These corresponded to partially shared internal segments. However, no significant similarity (BLASTn) was found between the flanking direct repeats when comparing GIs integrated at different t(m)DNAs.

### The *K. pneumoniae* GIs encode a variety of functions including those conferring virulence or drug resistance

To gain insights into the functions of the 3,540 gene families encoded in our set of GIs and prophages, they were annotated using Prokka [33] and the encoded proteins were mapped to the eggNOG v5.0 database and categorized by function using the eggNOG-mapper tool [34,35] (Figure 2H). In this analysis, only 1,305 proteins could be assigned to an eggNOG functional category, where 679 had predicted functions relating to information storage and processing, mainly in replication, recombination, repair, and transcription. Of note, a small proportion of proteins (292) were related to metabolism, with functions predominantly related to amino acid transport and conversion. The unclassified proteins were further analysed using customized annotation pipelines to identify possible overlooked predicted functions. We subsequently identified phage structural proteins, toxin-antitoxin systems, ICE-related proteins, and anti-restriction systems (Figure 2H). Also, we found a total of 286 genes distributed across 28 non-redundant GIs related to fimbriae assembly, type IV and type VI secretion systems, colibactin toxin and yersiniabactin and salmochelin siderophore production, drug efflux systems, as well as genes involved in iron, zinc, and manganese uptake and metabolism (Supplementary Table 4). Thus, a variety of known virulence and drug resistance determinants are encoded in GIs, raising the question regarding the role in these processes of the vast array of proteins with unknown function also located in this kind of mobile elements.

### Usage of tDNAs as integration sites for genomic islands and prophages

Next, we evaluated the usage of different t(m)DNAs as integration sites. From the 87 t(m)DNAs composing the core set, only 20 had a GI integration in at least one strain and with very dissimilar usage frequencies (Figure 3A). *ssrA* and five tDNAs (*arg3A, asn1D, phe1A, thr2A*, and *leu5A*) were occupied by GIs in more than 50% of the strains, outstanding *leu5A* harbouring a GI in 64 out of 66 strains (usage frequency=97%). Conversely, *phe1B* harboured a GI in only one strain and *ser2A* in two strains (usage frequencies of 1.5% and 3%, respectively). Additionally, the usage disparity among identical tDNAs located in different contexts reported previously for *asn1* tDNAs [8] was also observed for the 7 *met1* copies, where only *met1C* and *met1E* had integrated GIs. Further, while *phe1B* was largely not used, its paralog *phe1A* was used in more than 70% of the strains (Figure 3B). Concerning the diversity of islands found integrated into distinct tDNAs, the largest variety was observed for *leu5A* (33 different GIs), followed by *ssrA* and *thr2A* (22 and 21 different GIs, respectively). This demonstrates the remarkable dynamics of the *leu5A* locus and other specific t(m)DNAs, frequently associated with a very diverse set of GIs.

**Figure 3.**
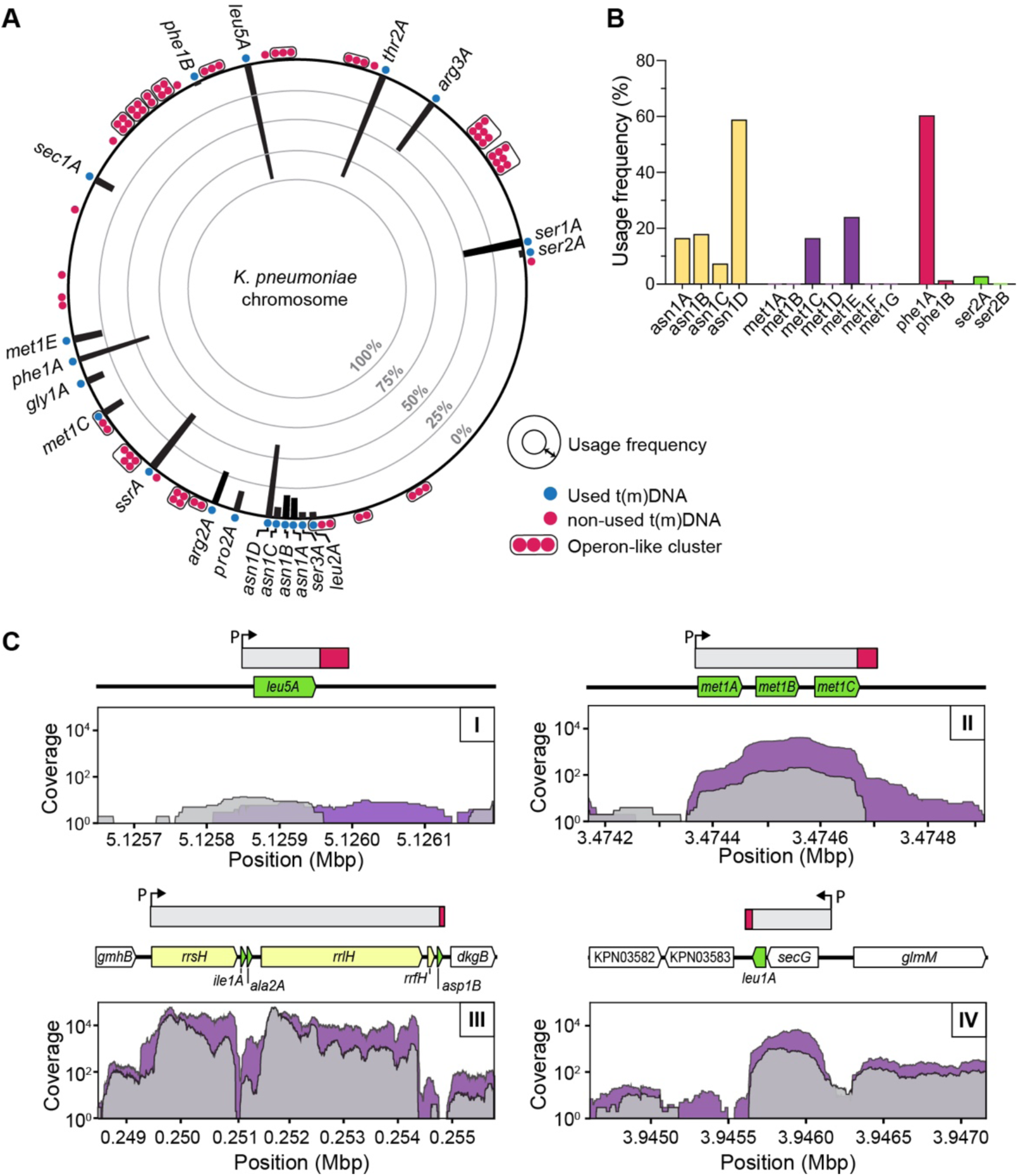
Usage of t(m)DNAs as integration sites for genomic islands in KpSC. (A) Radial plot showing the usage frequency of each t(m)DNA as integration sites (black bars pointing inside, according to the radial scale shown), as well as their approximate position in a representative KpSC chromosome. t(m)DNAs found associated with a GI in at least one strain are shown as blue circles, while non-used tDNAs are colored in red. t(m)DNAs forming part of an operon-like cluster are encircled together. (C) Structure of the transcriptional units comprising the t(m)DNAs from the core set, as approached by mapping previously published RNAseq data to the chromosome of the MGH78578 strain [40], as well as through bioinformatic prediction of promoters (P) and terminators (red boxes). Gray and purple areas show the coverage over chromosomal regions comprising t(m)DNAs (arrows in green) found transcribed alone (I), co-transcribed with other tDNAs (II), co-transcribed with rRNAs (arrows in yellow) (III), or co-transcribed with protein-coding genes (arrows in white) (IV).

Some features were identified amongst the genomic context of tDNAs that were used as integration sites. First, the locations of tDNAs are widely spread across the chromosome, except for one particular region where eight used tDNAs are located (*leu2A, ser3A, asn1A-D, pro2A*, and *arg2A*). Also, none of the tDNAs that form operon-like clusters nor those located inside rRNA operons were used as integration sites (Figure 3A and Supplementary Figure 3). One possible explanation is that selection against GI integration into t(m)DNAs forming polycistronic units would avoid disrupting the expression of co-transcribed downstream genes, as a consequence of separating them from their promoter. To test this hypothesis, we approached the structure of the transcriptional units (TUs) comprising each t(m)DNA through combining promoter and terminator prediction using the tools bTSSfinder [36], ARNold [37] and RhoTermPredictor [38], together with RNAseq-based transcriptional unit delimitation using the tool rSeqTU [39] and datasets previously generated for *K. pneumoniae* MG78578 [40] (Supplementary Table 5). We found t(m)DNAs that were part of monocistronic transcriptional units (type I); tDNAs co-transcribed with other tDNAs (type II); co-transcribed with rRNAs (type III); or co-transcribed with protein-coding genes (type IV) (Figure 3C and Supplementary Table 6). As expected, our analyses indicated that 18 of the 20 t(m)DNAs used as integration sites are transcribed as monocistronic units (type I). Chi-squared test supported the association between the usage frequency of the t(m)DNAs and the types of transcriptional units where they are located (p<0.05). Notably, the two exceptions were *met1C* and *leu2A*; both of these are co-transcribed along with other type-II tDNAs but constitute the last gene of the operon. All this evidence indicates that the genomic context influences the integration of GIs into tDNAs, avoiding the interruption of transcriptional units where one or more tDNA are co-transcribed with downstream genes.

Other possible factors explaining the asymmetry in the usage of different t(m)DNAs were explored. However, no association was observed between the usage frequency and 1) the conservation of the tDNA, 2) the conservation of the upstream and downstream contexts, and c) the %AT of the upstream region (Supplementary Figure 7A-D). Moreover, analysis using the MEME suite [41] showed no sequence motifs characteristic of the frequently used tDNAs or their contexts compared to those not used. Also, we hypothesized that transcriptional activity in the vicinity of a given tDNA could influence the integration of GIs. To test this, we exploited RNAseq data from *K. pneumoniae* MGH78578 [40] to calculate the mean expression level of the three genes immediately upstream and downstream of each t(m)DNA. We found no correlation between the frequency of GI insertion and the transcriptional activity of the region upstream or downstream of the t(m)DNAs (Supplementary Figure 7E-F). Factors affecting DNA topology including the nucleoid-associated proteins H-NS, Fis, and IHF may modulate accessibility to the t(m)DNA region and thus affect the integration of GIs. To test this hypothesis, we first corroborated that these three proteins from *K. pneumoniae* MGH78578 share over 94% identity with the respective proteins of *E. coli* K12 (Supplementary Figure 8A) and thus expecting they recognize highly identical binding sites. Then, using the PWMScan tool [42] and the binding sites matrices of H-NS, Fis, and IHF, experimentally determined for *E. coli* K12 available in the PRODORIC database [43] (Supplementary Figure 8B), we determined the abundance of predicted binding sites for these proteins in the 4-kbp regions upstream and downstream of each t(m)DNA. A slight significant positive correlation was only observed between t(m)DNA usage frequency and the number of downstream H-NS binding sites, while no correlation was registered for the other cases (Supplementary Figure 8C). Taken together, these results suggest that none of the factors tested, except avoiding TU disruption, could explain the frequency disparity in the usage of t(m)DNAs as integration sites.

### The role of the GI-encoded integrase in the selection of the integration site and GI evolution

To study the phylogenetic relationships among the integrases encoded in different GIs and possible correlations with their preference for specific tDNAs as targets for GI integration, a total of 125 integrase-coding sequences were extracted, translated, and aligned with MUSCLE [44] before inferring a maximum-likelihood phylogenetic tree with RAxML [45] (Figure 4A). Overall, the integrase proteins were highly diverse as evidenced by the deep branching of the tree, although all of them belonged to the tyrosine recombinase family, and clustered into three main clades (I-III). Inside these clades, we found smaller groups sharing conserved domains. Most of the integrases from clade I shared a phage P4-like catalytic C-terminal domain and an Arm DNA-binding N-terminal domain, while most from clade II harboured a shufflon-specific (phage Hp1-like) catalytic C-terminal domain. Some of these integrases also have an Arm or a SAM-like DNA-binding domain at their N-terminus. Lastly, clade III comprised mostly integrases with an uncharacterized site-specific tyrosine recombinase catalytic domain. Overall, similar integrases tended to target the same integration site, although some mixed clades were observed (Figure 4A). For instance, the subclade I.a, formed by 24 of the 28 integrases targeting *leu5* and all the integrases targeting *asn1*, comprised the integrase from arg2A-IV which shares 69% identity leu5A-XVI integrase and 57% identity with asn1-II, but only 45% with arg2A-XII integrase also located in this clade, and up to 33% identity with the other integrases from *arg2* GIs. Additionally, the integrase from phe1A-IV clustered between one of the encoded in asn1-XVII (58% identity) and leu5A-XXI integrase (56% identity). Also, subclade I.b comprised most of the integrases targeting *ssrA*, while subclade I.c grouped integrases targeting *thr2, arg3, sec1A*, and *arg2*; and I.d mainly integrases targeting the 5’ end of arg2A. Remarkably, we found that in strains 616 and A708 there were two GIs integrated at the *arg2A* locus, one at each end of this tDNA. No significant similarity (BLASTn) was found when comparing GIs harbouring integrases clustering close together but targeting different tDNAs (e.g. arg2A-IV and leu5A-XVI), arguing against a related origin of these elements. Additionally, PHASTER analysis indicated that although prophage integrases were found mostly associated with *met1*E, *arg2A*, and *arg3A* loci, they formed several mixed clades with GI integrases (for example leu5A-XX, ssrA-II, ssrA-IX, met1E-V, and sec1A-III) (Figure 4A). Moreover, we found 100% identical repeats from GIs and prophages targeting the same integration site (e.g. leu5A-XX and leu5A-XXIV, or arg2A-VI and arg2A-XI). This supports a common origin between some GIs and prophages, for instance, the latter progressively losing genes related to the viral particle assembly and acquiring genes relating to other functions.

**Figure 4.**
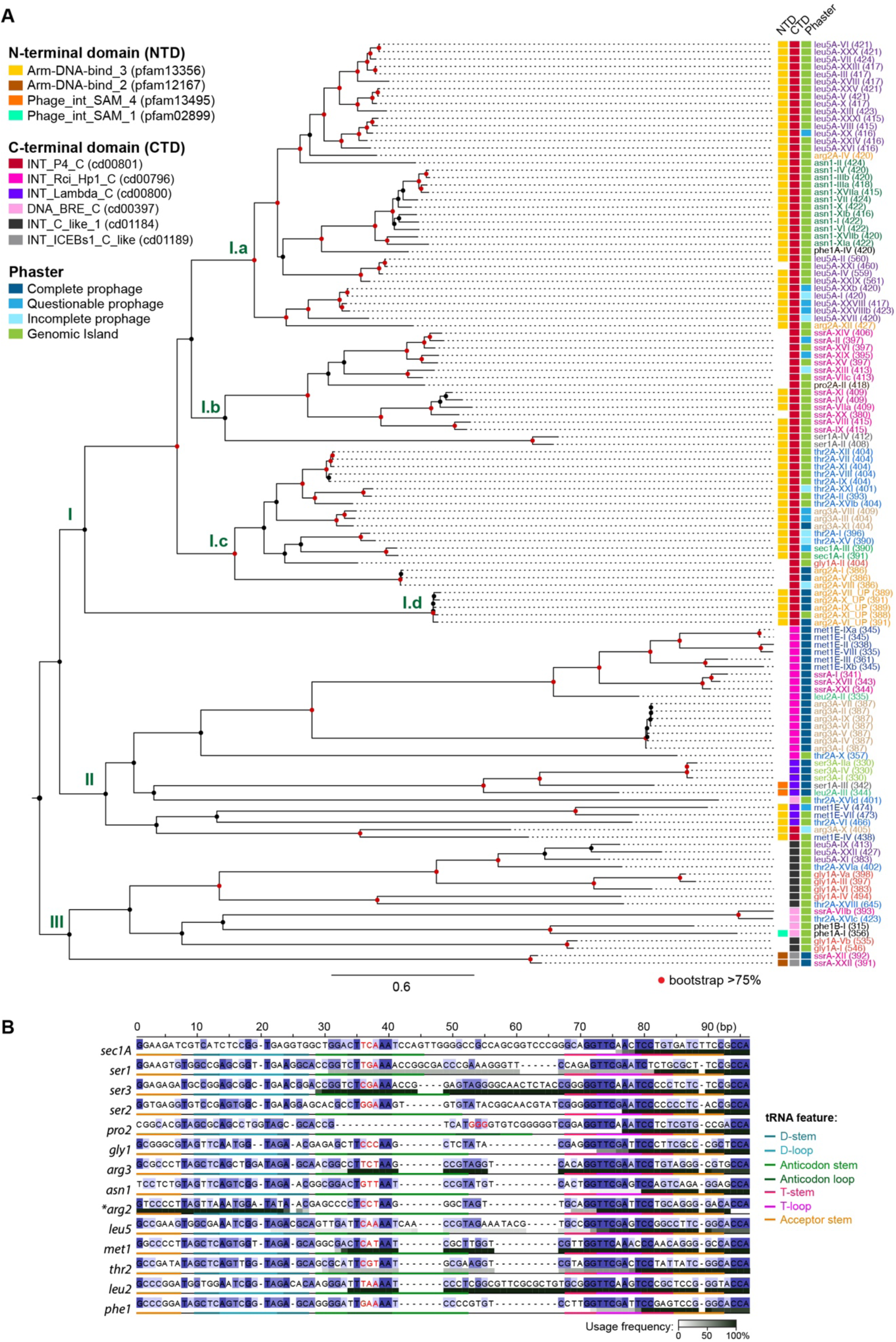
Diversity of KpSC GI-encoded integrases and t(m)DNA usage preferences. (A) The protein sequence of a total of 125 integrases identified among the non-redundant GI collection were aligned using MUSCLE, and this alignment was used to infer a maximum-likelihood phylogenetic tree. The displayed names correspond to the GI where the integrase is encoded, indicating in parentheses the total amino acid number. Shared name colors indicate integrases from islands integrated at the same tDNA. The numbers next to the nodes indicate the six main clades identified. (B) MUSCLE multiple alignment of tDNAs acting as integration sites. Coloring in blue indicates sequence conservation calculated from the alignment. Grayscale shading below each sequence indicates the frequency in which each base of the tDNA was used as a recombination site becoming part of the repeat, among our GI collection. Colored tDNA segments correspond to the key structural regions of the encoded tRNA. Bases corresponding to the anticodon are colored in red.

We next investigated the recombination site used by the integrases at different tDNAs, through identifying (when possible) the portion of these genes that formed part of the direct repeats. Additionally, we aligned the tDNAs found used as integration sites to identify possible shared sequence elements across their characteristic regions. Most of the recombination sites corresponded to the 3’ half of the tDNA including the CCA end, the 3’ half of the acceptor stem, the T-stem, and the T-loop (Figure 4B). In the case of *ser3, arg3, met1*, and *leu2*, the recombination site tended to be longer, including also the anticodon stem and the anticodon loop. As mentioned, we identified GIs integrated at the 5’ end of *arg2*, where the recombination site included the 5’ half of the acceptor stem, the D-stem, and the D-loop. Regarding *ssrA*, the repeats of their associated GIs corresponded mainly to its last 36 bases (tRNA-like domain and CCA end). Although the recombination sites at different tDNAs shared some bases, no correlation could be established between their sequence and the pattern of intermingled integrase clustering. Taken together, our results support that specific integrases from GIs and prophages evolved to recognize definite portions of a defined t(m)DNA, with no evidence of cross-integration. However, it remains unclear if these proteins could influence the preferential integration into one of the several copies of a given tDNA.

### Association of GIs with species from KpSC and evidence of GI acquisition through horizontal gene transfer

We aimed to investigate possible associations between specific GIs and certain species and lineages from KpSC, as well as to search for evidence of GI dissemination through horizontal gene transfer. For this, we used a core genome MLST approach to infer a maximum likelihood tree showing the phylogenetic relationships among the 66 KpSC strains included in this study using Roary and RAxML. A total of 3,564 non-redundant protein-coding genes present in >95% of the strains (95% protein identity cutoff) were included. For each strain, we depicted all the t(m)DNA-associated GIs present in its chromosome (GI profile), as well as its sequence type according to the current MLST scheme (https://bigsdb.pasteur.fr/) [19,46] (Figure 5). In general, although the same t(m)DNA integration sites are used across different lineages, variable GIs were found at these sites. Moreover, several GIs were found mainly associated with distinct *K. pneumoniae* lineages. For instance, as elements highly associated with clonal group 258 including the globally disseminated KPC-producing ST258 [10], we found asn1D-II and phe1A-I GIs (encoding DNA modification enzymes and many hypothetical proteins), as well as prophages arg3A-I, leu5A-I, and thr2A-I. Regarding hypervirulent clonal group 23, among the associated GIs we found leu5A-II, an 80-kbp potentially conjugative GI encoding a fimbrial operon and many helicases and nucleases, as well as the previously described islands GI-E492 (asn1C-VII) [8], ICE*Kp10* also known as GI-I (asn1D-III) [9,47], and ICE*Kp*1 (asn1D-IV) [15], each encoding factors related to virulence e.g. yersiniabactin, salmochelin, colibactin, the regulator of mucoid phenotype *rmpA*, and microcin E492.

**Figure 5.**
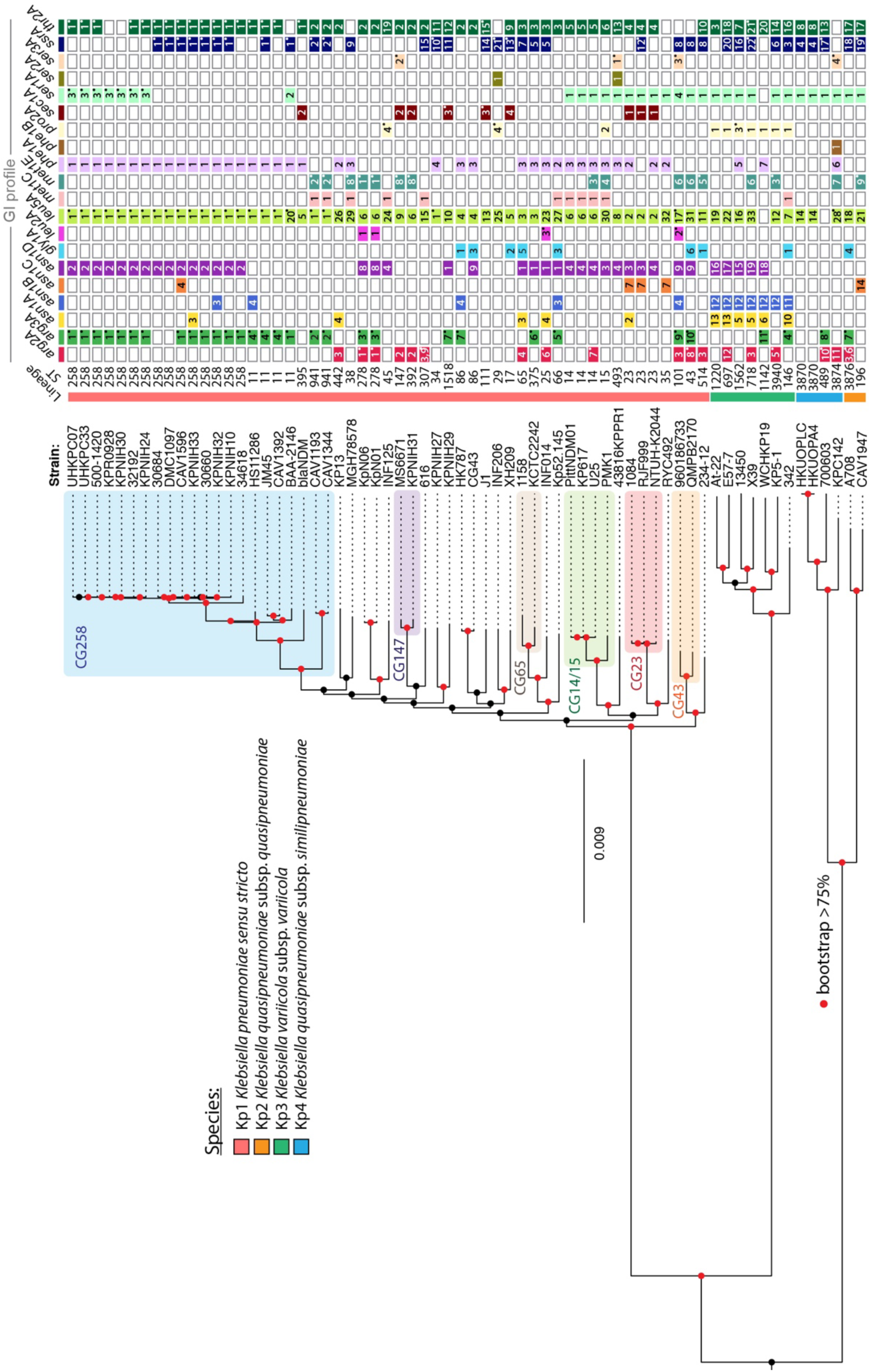
Phylogenetic inference among distinct KpSC strains supports GI dissemination through horizontal gene transfer. Maximum likelihood phylogenetic tree inferred from the concatenated alignment of 3,564 protein-coding genes shared by >95% of the strains. For each strain, the species, sequence type (ST), and the GIs integrated at each tDNA (GI profile) are indicated. Relevant clonal groups are shaded in colors. For the GI profile, shared colors indicate GIs integrated into the same tDNA, while the numbers indicate different GIs, as described in Supplementary Table 2. In strains 616 and A708, *arg2A* locus had 2 integrated GIs (one upstream and another downstream the tDNA). Boxes including a dot in the upper-right corner indicate elements identified as prophages by PHASTER analysis.

When inspecting in detail and comparing highly clonal strains, differences in GI content indicated acquisition or loss of GIs through horizontal gene transfer. Moreover, GIs that appear with relatively low frequency but spread across different and distant KpSC lineages would account for independent acquisition events. In this regard, despite the more recent shared ancestry amongst *K. pneumoniae* ST258 strains, we observed variation in GI content, including the loss of arg3A-I in strains 30684 and DMC1097, the loss of thr2A-I in KPR0928 and KPNIH30, as well as the acquisition of asn1C-IV in CAV1596, asn1A-III in KPNIH33 and asn1B-III in KPNIH32. Additionally, arg3A-VII, asn1B-IV, asn1D-IX, gly1A-I, and gly1A-III were variably present in the two highly related strains from ST86. Moreover, met1C-I and gly1A-I were sparsely present in strains from different *K. pneumoniae* STs and also in *K. variicola* ST146. Also, arg2A-III was found in some *K. pneumoniae* strains from ST442, 307, 101, and 514, but also in *K. variicola* X39 (ST718) and *K. quasipneumoniae* A708. All these results support a large diversity of GIs promoting chromosome variability, many of them being lost and/or disseminated through horizontal gene transfer.

### Conservation of tDNAs sequence, context, and usage as integration sites among different members of *Enterobacteriaceae*

Our global analysis of the *K. pneumoniae* t(m)DNAs indicates that both these genes and their upstream contexts are highly conserved among the four most common members of KpSC (Kp1-Kp4). To explore the degree of conservation of these elements among other members of *Enterobacteriaceae*, we identified and compared the asparagine tDNAs from the fully assembled chromosomes of 13 strains of *Yersinia pestis*, 6 *E. coli*, 9 *Shigella flexneri*, 11 *Salmonella enterica*, 3 *Salmonella bongori*, 7 *Serratia marcescens*, and 7 *Enterobacter cloacae* (Supplementary Table 7). We found in all these species, as in KpSC, 3 to 4 copies of the *asn1* tDNA (GUU anticodon). Remarkably, the *asn1* tDNAs showed 100% identity across all the strains analysed, except for strain *S. flexneri* 2a301, where 2 out of 4 *asn1* genes were identical to those in the other species, while the third showed a single nucleotide substitution and the fourth harboured a 1-bp insertion. In all the other *S. flexneri* strains, 100% identity was observed across the 4 *asn1* genes. Additionally, the immediate upstream contexts of the *asn1* tDNAs found in KpSC were also conserved in most of the species, although additional variable contexts were observed (named *asn1F* to *asn1J*) (Figure 6A). The high conservation of tDNAs among these bacterial species implies that they would offer equivalent integration sites in their chromosome, potentially enabling their sharing and spreading through inter-species horizontal transfer. As done with *K. pneumoniae* strains, upon the identification of the virgin context of the *asn1* tDNAs, we were able to search for GIs integrated within these sites for each of the other species (Supplementary Table 8). A total of 32 GIs with sizes ranging from 2.8 kbp to 173 kbp were identified, 22 of them harbouring both direct repeats and an integrase-coding gene, 3 only the integrase, 1 only the repeats, and 6 lacked these features. Some shared content was found with KpSC asn1-associated GIs, outstanding asn1A GIs from 4 *E. coli* and 6 *Y. pestis* strains corresponding to members of the ICEKp family. Among the functions encoded, we found the yersiniabactin siderophore and other iron uptake systems, multidrug efflux pumps, proteins related to attachment and invasion, type-IV secretion systems, and flagella assembly.

**Figure 6.**
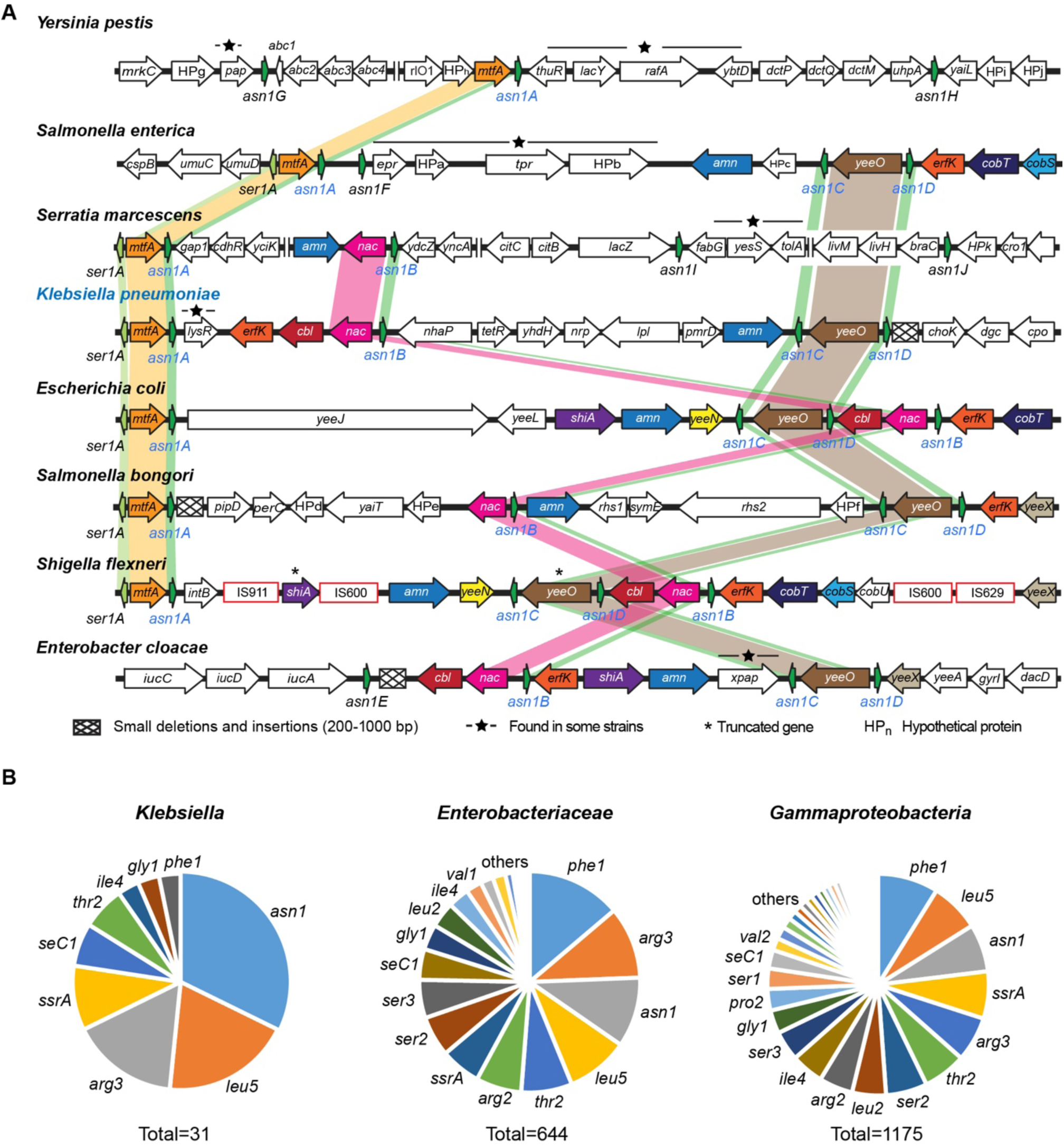
The genomic contexts of the *asn1* tDNAs as well as the t(m)DNA usage patterns are conserved among different species of *Enterobacteriaceae*. (A) Scheme showing the chromosome region comprising the 3-4 copies of *asn1* asparagine tDNAs from eight *Enterobacteriaceae* species, named *asn1A* to *asn1D* as for the tDNAs found in KpSC, or *asn1E* to *asn1J* as additional genomic contexts were identified among the other species, following the nomenclature used in this study. Colored arrows indicate conserved genes across chromosomes from different species. Colored connecting ribbons indicate tDNAs and their immediate upstream contexts which are conserved. (B) Frequency in which different t(m)DNAs were used as integration sites in *Klebsiella, Enterobacteriaceae*, and Gammaproteobacteria, according to the genomic islands present in the Islander database.

To gain insights regarding the preferential usage of a specific set of tDNAs as integration sites for GIs in other bacteria, we downloaded the Islander database [48], currently comprising 4065 islands identified among 1313 archaeal and bacterial genomes (31 of these from *K. pneumoniae*). After classifying these GIs by the reported integration site and the species of origin, we observed that *leu5, thr2, asn1, phe1*, and *ssrA* (the top 5 most used in KpSC according to our analyses) were also the most frequent integration sites (top 6) among all the GIs present in the Islander database from *Enterobacteriaceae* and more distant members within Gammaproteobacteria (Figure 6B). Thus, the t(m)DNAs and their contexts remained conserved through thousands of years of evolutionary history, where a subset of them was selected to act as hotspots of genetic variability. This constraint for a specific set of tDNAs as integration sites would have evolved before the divergence of Gammaproteobacteria, and probably results of interacting forces driving bacterial chromosome evolution towards allowing the integration of GIs, preserving the tDNAs and their genomic contexts, and the evolution of integrases to preferentially target monocistronic tDNAs.

## DISCUSSION

Accumulating evidence from the analysis of thousands of genomes indicates that the contribution of horizontal gene transfer to bacterial chromosome evolution has been largely underestimated [49,50]. This would be particularly true in bacteria living in highly diverse and populated microbial communities such as the gut microbiome, which is also an entrance route for a variety of pathogenic strains, many of them from *Enterobacteriaceae* [51,52]. In this context, mobile genetic elements are key drivers of pathogen evolution mediating the exchange and dissemination of genes involved in virulence, antibiotic resistance, and fitness [53]. This is the case of *K. pneumoniae*, a priority pathogen showing the rapid development of multidrug-resistant and hypervirulent strains, and remarkable genomic plasticity [18,54]. Here, we provided evidence that t(m)DNAs are key loci mediating chromosomal diversity in this species, acting as integration sites for an ample repertoire of GIs and prophages, many of them encoding factors potentially involved in virulence and drug resistance.

tDNAs and *ssrA* are very particular genes with various associated properties and functions [55]. Although studied for a long time, an outstanding aspect that remains obscure is why these genes, normally constituting <2% of the bacterial genome, are used as integration sites more frequently than other genes (i.e. protein-coding genes). Three non-mutually exclusive hypotheses have been proposed in this regard [56]. The first one is based on selection, postulating that although new recombination sites could arise at any chromosomal locus, t(m)DNAs would better fit the biology of GIs and prophages. In this context, one determining factor would be their low variation, showing a divergence rate per base pair 4- to 9-fold lower than for protein-coding genes, and thus offering more stable recombination sites in a broader host range [56–58]. Indeed, we found high conservation of both t(m)DNAs and also their upstream genomic contexts among KpSC and representatives from other *Enterobacteriaceae*. On the other hand, it was proposed that the 3’-end of mature tRNAs could hybridize to one strand of the encoding tDNA, and the hybrid structure might somehow improve integrase recognition and action [59]. However, we found a specific set of integrases targeting the 5’ end of *arg2*, as well as two GIs with a repeat lacking the CCA end, arguing against the 3’ end and the CCA sequence as general requirements for the recombination sites. In a second hypothesis (drift), sequences similar to an established t(m)DNA recombination site (*attB*) are most likely to be found in another t(m)DNA, thus, a few mutations in the *attP* site can create a match to a new t(m)DNA gene [56]. In this line, *ssrA* was shown to contain a CCA 3’-end and an acceptor stem that are identical to those of alanine tDNA and indeed the tmRNA is aminoacylated with alanine [59]. However, our results indicate that even tDNAs recognized by more similar integrases offer *attB* sequences with limited similarity (e.g *leu5* and *asn1*). Further, while *ssrA* had an integrated GI in more than 50% of the strains, no GIs were found integrated at any of the *ala1* and *ala2* tDNAs. Thus, we were unable to deduce possible drifts from one integration site to another.

The third hypothesis proposes that integrases recognize some features shared by the t(m)DNAs that are absent in other kinds of genes, for instance, characteristic segments with sequence symmetry and complementarity. This is supported by bioinformatic evidence indicating that, for most of the t(m)DNAs, recombination occurs at specific sublocations inside these genes, mainly two with flanking symmetry (anticodon-loop and T-loop) and a third one at the asymmetric 3’ end. Further, different integrase subfamilies would use either the symmetric sublocations or the asymmetric sublocation exclusively, although different t(m)DNAs can be used within any subfamily [56]. Experimental evidence supporting this sublocation specificity was provided for the integrase of the ICE*Afe1* conjugative GI from *Acidithiobacillus ferrooxidans*, where 19 bp encompassing the anticodon loop but not the rest of the tDNA sequence were required for the *in vivo* integration in a plasmid fusion assay [60]. Overall, our results support integrase subfamilies targeting distinctive sublocations. For instance, in most of the integrases from group I.a targeting *leu5, arg2, asn1*, and *phe1*, the *attB* site started in the T-loop. Conversely, integrases targeting some of these tDNAs but clustering outside group I.a, e.g. leu5A-IX or arg2A-VII, showed a different and novel sublocation preference: The D-loop. Notably, arg2A-VII and other GIs harbouring an integrase from clade I.d integrated upstream of *arg2*, in contrast to all other the GIs integrating downstream the target DNAs. These integrases carried the same N- and C-terminal domains than most of the integrases from the group I, indicating that sublocation preferences would be defined at a smaller scale than domain content. Otherwise, for most integrases of group II, the *attB* extended from the anticodon loop to the CCA end. This evidence supports sublocation preferences as a relevant force directing integrase and site recognition evolution.

Beyond hypotheses explaining preferential t(m)DNA usage as integration sites over other types of loci, until now no clues were provided regarding possible factors driving a constraint for the usage of a subset of them. In this regard, we found only 20 out of 87 t(m)DNAs with integrated GIs among our strain collection. Some of this same set of t(m)DNAs were shown as integration hotspots in other bacteria [14,56,59,61], suggesting evolutionary conservation of this bias. Our evidence indicated that most of the t(m)DNAs used as integration sites would be part of monocistronic TUs, being the only two exceptions (*met1C* and *leu2A*) tDNAs located at the end of the operon. Thus, purifying selection would be acting against the disruption of transcriptional units upon GI integration. In this scenario, GIs could integrate into other tDNAs but the increased relative fitness costs of disrupting a polycistronic unit would result in the rapid purging of such strains from the population, constraining the evolution of integrase-*attB* recognition to t(m)DNAs where such disruption and cost do not occur. Despite the above, although *asp1A* and *ser2B* are also predicted as monocistronic units we did not find GIs integrated into these tDNAs. Moreover, even when forming part of single TUs and having a 100% sequence identity, some tDNAs located in different positions of the chromosome were used with very dissimilar frequencies. This was also evidenced upon analysis of the ICE*Kp* family and GI-E492, both *asn1*-associated GIs, where marked preferences for one of the four copies of *asn1* were detected [8,9]. Hence, further studies are required to identify additional factors behind usage bias towards specific t(m)DNAs.

Our analysis of 66 *K. pneumoniae* chromosomes allowed the identification of 162 non-redundant GIs, contributing to a significant increase in the known islands from this species, and yet suggest that a considerable number of *K. pneumoniae* GIs remain undiscovered. The identified GIs encoded *∼*28% of the chromosomal accessory genome, supporting a major role of GIs mediating chromosome diversity, although it is clear that a large proportion (∼72%) of chromosomal accessory genes are not part of t(m)DNA-associated elements, and warrant further investigation. This additional accessory genome could include GIs integrated at other loci, other kinds of mobile elements such as transposons or integrons, or can be originated from rearrangements mediated by homologous recombination e.g. as described for the *Klebsiella* capsule (K) and lipopolysaccharide (O) loci [62,63], which are estimated to account for ∼11% of the total *K. pneumoniae* pan-genome [54].

Through comparing core-genome phylogeny with GI profile, we provided evidence for the mobilization and horizontal transfer of several GIs in diverse *K. pneumoniae* lineages. In this regard, previous work experimentally demonstrated the excision of some *K. pneumoniae* GIs [8,15] and the transfer by conjugation of the ICE*Kp1* element, a well-studied conjugative island from *K. pneumoniae* [15]. Moreover, we previously showed that some *asn1*-associated GIs including GI-E492 harbour a cryptic transfer origin similar to that of ICE*Kp* elements and thus could be mobilizable through conjugation, a property shared by a poorly studied group of genetic elements named mobilizable genomic islands [14,64]. Seven out of 19 GIs targeting *asn1* described here harboured this oriT, and it was found only in *asn1*-associated GIs. Furthermore, our analysis of additional bacterial species revealed that *asn1* and likely other t(m)DNAs are conserved across *Enterobacteriaceae*, and that a similar subset of t(m)DNAs used as integration sites in KpSC would be also the preferred ones across this family and also Gammaproteobacteria. This suggests that KpSC GIs would find the same integration sites in different species of this family, potentially allowing their inter-species horizontal spread. However, in opposition to this idea, although many of the ICE*Kp* elements could disseminate by conjugation, analysis of their presence among *∼*2500 KpSC genomes and 11 additional species from *Enterobacteriaceae* indicated that the vast majority of these elements (97%) were found in *K. pneumoniae* [9]. Moreover, only 2 different GIs found in other species showed significant sequence similarity and coverage with the KpSC GIs identified. This suggests that some barriers would act against inter-species GI dissemination (e.g. restriction-modification or CRISPR-Cas systems).

Several questions remain to be addressed regarding t(m)DNAs and their role as relevant *loci* mediating chromosome diversity including what additional factors would influence integration site usage? Is the usage bias maintained across a broader sample of *K. pneumoniae* genomes and/or in other species? Can usage bias be observed directly in experimental integration assays? What are possible barriers to inter-species GI dissemination? What is the functional relevance of many GIs encoding mostly hypothetical proteins? Future studies are required to shed light in these directions.

## CONCLUSIONS

The systematic comparison of the t(m)DNAs across different strains allowed us to determine the typical set of these genes present in highly conserved regions of the KpSC chromosome. The knowledge of each t(m)DNA region in its virgin configuration was leveraged to discover and characterize an unprecedented number of highly diverse KpSC GIs and prophages, several of them likely being disseminated by horizontal gene transfer. This mobility is mediated by a wide repertoire of integrases targeting specific sublocations at specific t(m)DNAs and by type-IV secretion systems and/or an oriT for conjugation. Remarkably, we found a strong bias for the usage of a subset of the tDNAs as integration sites, likely towards avoiding disruption of polycistronic transcriptional units. This raised important questions and clues regarding the fundamental properties of t(m)DNAs as integration sites for mobile genetic elements and drivers of genome evolution and pathogen emergence.

## METHODS

### Source of DNA sequences

The genome sequences analysed in this work were retrieved from the National Center for Biotechnology Information (NCBI) database. The accession numbers and relevant metadata of KpSC strains and strains from other *Enterobacteriaceae* species are provided in Supplementary Table 1 and Supplementary Table 7, respectively.

### Identification of tDNAs and classification according to their conserved genomic contexts

The primary identification of all the t(m)DNAs was performed using ARAGORN v1.2.38, determining the anticodon carried and the coordinates [20]. Then, each t(m)DNA was analysed to identify its genomic context, defined arbitrarily as the three protein-coding genes located upstream and downstream, considering both their nucleotide sequence and organization, and also the presence of adjacent tDNAs and rRNAs. Based on this, the tDNAs were annotated using the nomenclature system proposed in this work, which can be automatically performed using our tool Kintun-VLI. tDNAs characteristic regions were determined using the search tool of the Transfer RNA database (http://trna.bioinf.uni-leipzig.de/) [65].

### Evaluation of the sequence conservation among tDNAs and their contexts

Sequence identity calculations among t(m)DNAs and their flanking regions were carried out by multiple alignments using ClustalW v2.1 [22] and custom scripts developed using Biopython [66]. The identity values showed in Figure 1 correspond to the mean percentage identity among all the pairwise alignments of the tDNAs encoding the same anticodon and located in an equivalent genomic context, from a total of 5,624 t(m)DNAs identified in our dataset. For t(m)DNA-flanking region identity calculations, we used custom BioPython scripts to annotate all the t(m)DNA-associated GIs in each chromosome and then, if found, to remove these regions to reconstruct the virgin locus configuration. Afterward, the sequences corresponding to 4 kbp upstream and 4 kbp downstream from each t(m)DNA were extracted and used to perform multiple alignments. To calculate the identity based on the relative position from the t(m)DNA (both for upstream and downstream sequences), a 200 bp starting region was considered, determining the mean identity of pairwise alignments inside each group of sequences. Then, the next 200 bp of the flanking regions were added to the sequence, repeating the alignment and identity calculations until completing the total extension.

### Codon usage frequency calculation

To determine the codon usage preferences, we used Roary v3.11.2 [32] to extract 3,564 non-redundant protein-coding genes (95% protein identity cutoff) present in >95% of the 66 *K. pneumoniae* strains described in Supplementary Table 1. Then, we used GCUA v1.0 [67] and custom scripts to count all the codons present in a representative sample of these genes (part of the pangenome_reference.fa file generated by Roary) and calculate the codon usage frequency (F_cod_), defined as the count of a specific codon, divided by the total count of codons specifying the same amino acid. A similar approach was conducted with the genomes of 72 *E. coli* strains from the ECOR collection [68,69], along with 3 NCBI reference strains (K-12 substr. MG1655, O157:H7 str. Sakai, and DSM30080), extracting and counting the codons from a total of 3037 CDS.

### Identification of t(m)DNA associated genomic islands

GI identification was based on using BLAST and ProgressiveMauve [29] to evaluate the shift of the known conserved downstream context, according to the virgin locus configuration defined for each tDNA. Thus, a GI was defined as any DNA segment >2.5 kbp located between the 3’ end of a given tDNA and its characteristic conserved downstream sequence. The identified putative GIs were searched for direct repeats and integrase coding genes by using BLASTn and BLASTp, respectively. This GI identification pipeline was automated and implemented as part of Kintun-VLI. The correlation between the total chromosome length and either the number of GIs or the total GI length, as well as between the total number of CDSs and the GI-encoded CDSs were evaluated performing a linear regression and determining the Spearman’s correlation coefficient (r) with the associated P-value.

### Functional annotation of the genes encoded in genomic islands

The non-redundant GI collection was annotated utilizing PROKKA v1.13 [33]. Using Roary, a total of 3540 non-redundant protein families were identified, which were functionally categorized using the eggNOG_mapper v2 tool [35] and the eggNOG v5.0 database [34]. Tabulated data of the categorization was used to create the sunburst chart shown in Figure 4 using the sunburstR v2.1.1 package in R v3.6.0 [70]. Also, the predicted genes were further inspected searching for putative functions using BLASTp v2.6.0 [28] along with the VFDB [71] (downloaded on June 25, 2019) and CARD [72] (version 3.0.2, Augist, 2019) databases, and also using the tools available as part of the PATRIC Bacterial bioinformatics resource center (https://www.patricbrc.org/) [73]. To evaluate the results, matches with coverage greater than 80% and an identity greater than 90% were considered. Phage detection and classification were performed using PHASTER [30].

### Comparison of the nucleotide sequence and gene content among different genomic islands

The GI nucleotide sequences were compared performing pairwise alignments using BLASTn v2.6.0. Positive matches, defined by thresholds of >80% identity and >50% coverage of the smallest GI, were plotted as a heatmap using the ComplexHeatmap v2.1.0 package [74] in R v3.6.0. Diagrams showing shared regions between different GIs were constructed using EasyFig v2.2.2 [75].

The total number of gene families present in the collection of 162 non-redundant GIs were determined using Roary. Then, the gene presence/absence file from Roary output and the functional categorization performed using eggNOG_mapper v2 tool were used to plot as a heatmap the presence or absence of each gene family inside each genomic island (ordered by functional category), using the ComplexHeatmap v2.1.0 package.

### Prediction of the transcriptional units comprising t(m)DNAs

The prediction of the t(m)DNA transcriptional units was carried out by integrating bioinformatic promoter and terminator identification, together with mapping RNAseq data over these regions exploiting coverage fluctuations to define the approximate start and end of the units. Promoter prediction was performed with the bTSSfinder tool [36] and terminator sequences were predicted using ARNold [37], and RhoTermPredictor [38]. Predicted promoters and terminators were manually reviewed by comparing them with the promoters and terminators determined for equivalent regions in the chromosome of *E. coli* K12, as published in the EcoCyc repository (accessed in August 2019) [76]. Transcriptional unit delimitation through RNAseq mapping was made using previously published data from *K. pneumoniae* MGH78578 available from Sequence Read Archive (SRA), and the rSeqTU tool, a machine-learning-based R package for prediction of bacterial transcription units [39]. In total, 12 experiments were considered for the analysis (Supplementary Table 5), plotting for each type of transcriptional unit (Figure 3) the 2 experiments showing the highest overall coverage values, and to define the approximate transcription start and end sites of each region (Supplementary Table 6). Reads mapping against the MGH78578 chromosome was performed using Bowtie2 [77] and the coverage plots for the regions showed in Figure 3 were made by using the Matplotlib v3.1.1 library [78], DNA Feature Viewer v3.0.1 [79], and custom Python scripts.

### Construction of an accumulation curve of *K. pneumoniae* genomic islands

The GIs accumulation curve was calculated using the vegan package in R, as previously described (Holt, et al., 2015). To assess whether the number of new types tends to increase or converge to a constant value, the curve obtained was adjusted according to the Heaps’ law, as described by Tettelin et al., 2008, using R. Accordingly, the equation N(x) = kx^(1-α)^ was used, where N(x) is the number of new GIs identified as a function of the number of strains analysed (x), and (1-α) corresponds to the growth exponent, which can be used to determine if the set is open or closed. For α>1, the GI accumulation is considered closed, as its size approaches a constant as more genomes are sampled. Conversely, for α<1, the GI accumulation is an increasing and unbounded function of the number of genomes considered.

### Average GC content calculation for *K. pneumoniae* genomes

Total GC content was calculated for each of the 66 chromosomes using the Perl script get_gc_content.pl (https://github.com/mrmckain/Herbarium_Genomics/blob/master/get_gc_content.pl). The values obtained for each strain were used to calculate the mean and the standard deviation of the whole set.

### Construction of the integrase phylogenetic tree

The protein sequences of 125 integrases encoded in the non-redundant set of GIs were aligned using MUSCLE [44], and the resulting alignment was used to infer a maximum-likelihood tree using RAxML v8.2.12 [45] set to automatically determine the best substitution model for proteins (-m PROTGAMMAAUTO) and 100 bootstrap iterations. The calculation was performed a total of 5 times varying the seed parameters, and the tree with best likelihood values was selected and then mid-point rooted and plotted using FigTree v1.4.4 (https://github.com/rambaut/figtree/releases).

### *K. pneumoniae* species complex phylogeny

We used Roary to extract 3,564 non-redundant protein-coding genes present in >95% of the strains (95% protein identity cutoff). Then, the aligned core chromosomal genes were used to infer a maximum-likelihood phylogeny with RAxML using the GTR+Gamma substitution model (-m GTRGAMMA) and 100 bootstrap iterations. The calculation was performed a total of 5 times varying the seed parameters, and the best likelihood tree was mid-point rooted and plotted using FigTree.

### Analysis of the genomic islands present in the Islander database

The complete Islander database [48] (as available in May 2019) was downloaded, comprising a total of 4065 islands in 1313 archaeal and bacterial genomes. The information retrieved included the name of the GI, its length, its GC content, the lineage of the host, and the integration site. These data were included in dynamic tables and analysed using spreadsheet editors and GraphPad Prism 8.3.1, to calculate and plot the frequency of t(m)DNA usage as integration sites inside different taxonomic groups.

### Search for conserved sequence motifs and binding sites for nucleoid-associated proteins in the contexts of the t(m)DNAs

Possible conserved sequence motifs in the surroundings of the t(m)DNAs were evaluated using the MEME v5.1.1 tool available from the MEME Suite [41]. Two analysis modes were conducted using the *K. pneumoniae* MGH78578 genome as reference: The *Classic* mode to search for conserved motifs among a collection of sequences, and the *Differential Enrichment* mode to search for enriched motifs in a set of sequences compared to a control group of sequences where the motifs are expected to be absent. Using Biopython scripts, the sequence of the 4-kbp regions upstream and downstream of each t(m)DNAs (excluding sequences identified as genomic islands) were extracted and grouped/analysed as follows. Under the *Classic* mode, three groups were considered: (i) regions flanking the t(m)DNAs used as integration sites; (ii) regions flanking t(m)DNAs that are not used as integration sites; and (iii) regions flanking the whole set of t(m)DNAs. Under the *Differential Enrichment mode*, these groups and also sub-groups of t(m)DNAs used as integration sites were compared. Regarding the distribution of the expected motif, we used the “any number of repetitions” (*anr*) for the Classic analysis and option zero or one occurrence per sequence (zoops). All the other parameters were set as default.

Identification of the binding sites of nucleoid-associated proteins IHF, Fis, and H-NS was performed for the *K. pneumoniae* MGH78578 chromosome. For this, we retrieved from the PRODORIC database [43] the position weight matrices representing the consensus sequence of these binding sites defined for *E. coli* K12. Using these matrices and the tool PWMScan [42] we identified possible binding sites with a p-value<0.00001 across the MGH78578 chromosome. Then, we counted the number of predicted binding sites for each protein present in 4-kbp regions upstream and downstream of each t(m)DNA, testing the possible correlation between the number of sites and the usage frequency as integration sites.

## AUTHORS’ CONTRIBUTIONS

AM, CBP, KH, KW, ML, and RL conceived and designed the study. AM, CBP, KW, ML, MV, PA, and RA, conducted the computational work and data analysis. AM, CBP, PA, and RA participated in Kintun-VLI tool design, programming, and testing. AM, CBP, KW, ML, and RL wrote and substantively revised the manuscript. All authors read and approved the final manuscript.

## ACKNOWLEDGEMENTS

We want to acknowledge Carlos Serrano Pinto for its contribution to the prediction of the t(m)DNAs transcriptional units’ structure.

## ADDITIONAL FILES

**Supplementary Figure 1**. Description and scheme of the proposed nomenclature to name KpSC tDNAs based on the encoded anticodon and the conservation of their genomic contexts.

**Supplementary Figure 2**. Sequence conservation of the t(m)DNAs and their contexts among 66 KpSC chromosomes.

**Supplementary Figure 3**. Schematic representation of the virgin contexts in which are located the different tDNAs that compose the core set of KpSC.

**Supplementary Figure 4**. Diagram showing the presence or absence of the 3,540 GI-encoded non-redundant gene families across the different GIs identified in this study, categorized by function using the eggNOG database.

**Supplementary Figure 5**. Significant matches in pairwise sequence alignments among the GIs described in this study.

**Supplementary Figure 6**. Shared DNA regions among different t(m)DNA-associated GIs found in *Klebsiella* genomes.

**Supplementary Figure 7**. Evaluation of the possible correlation between the usage frequency of t(m)DNAs as integration sites and several properties of these genes and their genomic contexts.

**Supplementary Figure 8**. Evaluation of the possible correlation between the usage frequency of t(m)DNAs as integration sites and the abundance of H-NS, IHF, and Fis predicted binding sites in the upstream and downstream contexts.

**Supplementary Table 1. (spreadsheet)** List and properties of the *K. pneumoniae* strains whose genomes were analysed in this study.

**Supplementary Table 2. (spreadsheet)** Number and distribution of tDNA copies compared to codon usage in *K. pneumoniae* and *E. coli* genomes.

**Supplementary Table 3. (spreadsheet)** Classification and relevant features of the genomic islands and prophages integrated at t(m)DNAs in the set of 66 *K. pneumoniae* strains analysed in this study.

**Supplementary Table 4. (spreadsheet)** GI-encoded proteins related to virulence, drug resistance, and horizontal gene transfer.

**Supplementary Table 5**. RNAseq datasets used to predict transcriptional units comprising t(m)DNAs by means of the tool rSeqTU.

**Supplementary Table 6. (spreadsheet)** Predicted transcriptional units constituted by the tDNAs present in the *K. pneumoniae* MGH58578 chromosome.

**Supplementary Table 7. (spreadsheet)** Strains from different species of *Enterobacteriaceae* whose genomes were analysed in this study.

**Supplementary Table 8. (spreadsheet)** Asparagine tDNA usage frequency and associated GIs found in different strains of *Enterobacteriaceae*.

