## Supplementary material for "Properties of genes encoding transfer RNAs as integration sites for genomic islands and prophages in *Klebsiella pneumoniae*"

**-Supplementary Table 7. (spreadsheet)** Strains from different species of *Enterobacteriaceae* whose genomes were analysed in this study.

**-Supplementary Table 8. (spreadsheet)** Asparagine tDNA usage frequency and associated GIs found in different strains of *Enterobacteriaceae*.

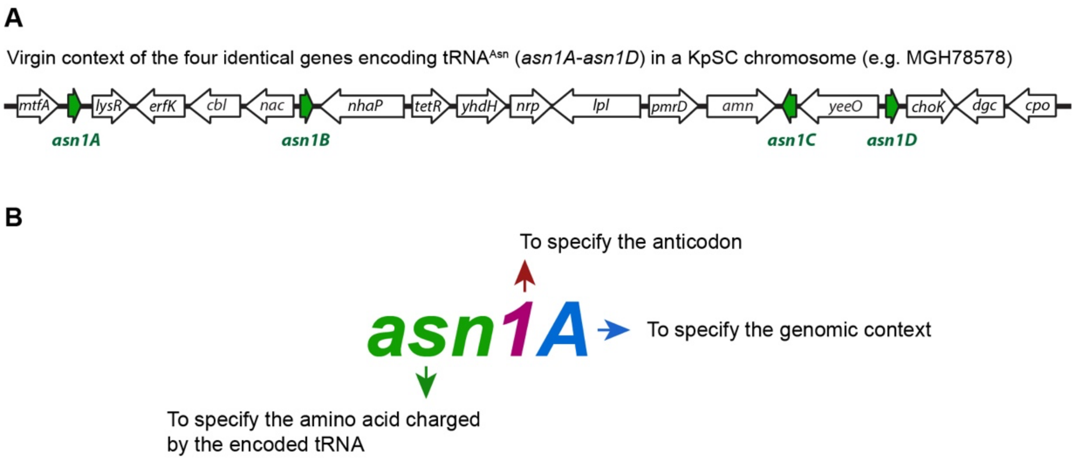


Nomenclature directions:

1) Following the convention for naming bacterial genes, all the letters should be used in italics.

2) The first lower-case letters correspond to the three-letter code of the amino acid charged by the encoded tRNA.

3) The following Arabic number indicates one of the possible anticodons to translate this specific amino acid. It must be asigned following the anticodon codes described in this study.

4) The following capital letter identify the conserved and characteristic genomic context of each tDNA. It must be asigned following the collection of genomic contexts described in this study.

**Supplementary Figure 1. Description and scheme of the proposed nomenclature to name the KpSC tDNAs based on the encoded anticodon and the conservation of their genomic contexts.** A: Genetic organization of the KpSC chromosome region comprising the four identical copies of the asparagine tDNA in absence of any integrated mobile element (virgin context). These tDNAs encode the same anticodon (GUU) but are located in different contexts (adjacent to different genes). Thus, following the directions shown in (B), they were named *asn1A*, *asn1B*, *asn1C*, and *asn1D*. The number associated with the anticodon and the letter associated with the context were defined arbitrarily and should be assigned following the code shown in the Supplementary Table 2 (anticodon nomenclature) and Supplementary Figure 3 (genomic context classification). Automated KpSC tDNA annotation using this nomenclature can be performed using our Kintun-VLI tool (https://github.com/GMI-Lab/Kintun-VLI).

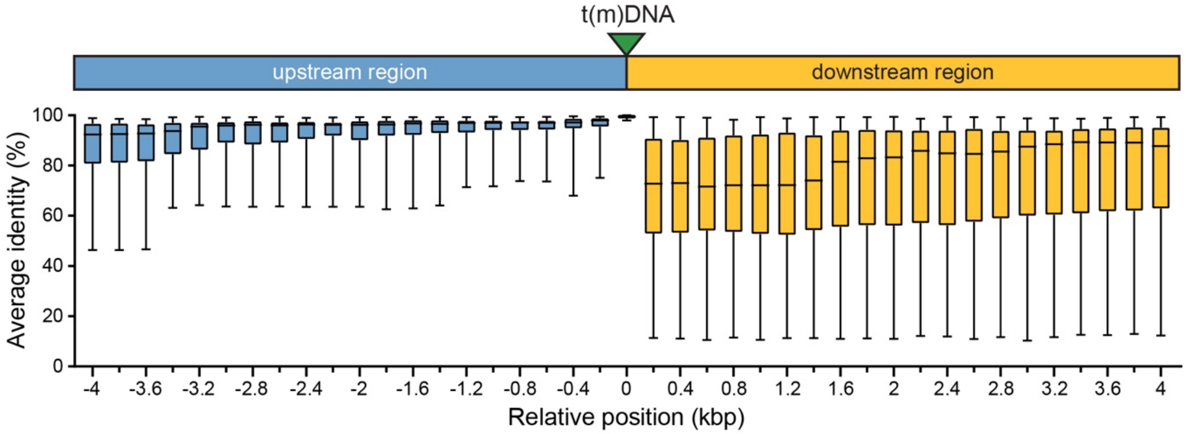


**Supplementary Figure 2. Sequence conservation of the t(m)DNAs and their contexts among 66 KpSC chromosomes.** Equivalent DNA segments comprising a defined t(m)DNA plus 4 kbp of the adjacent upstream and downstream regions were extracted from all the chromosomes and aligned using MUSCLE. From the alignment, the average identity was calculated in growing windows of 400 bp (starting from the borders of the t(m)DNA). The per-window average identity values observed for the contexts of all the tDNAs from the core set are shown as a box plot.

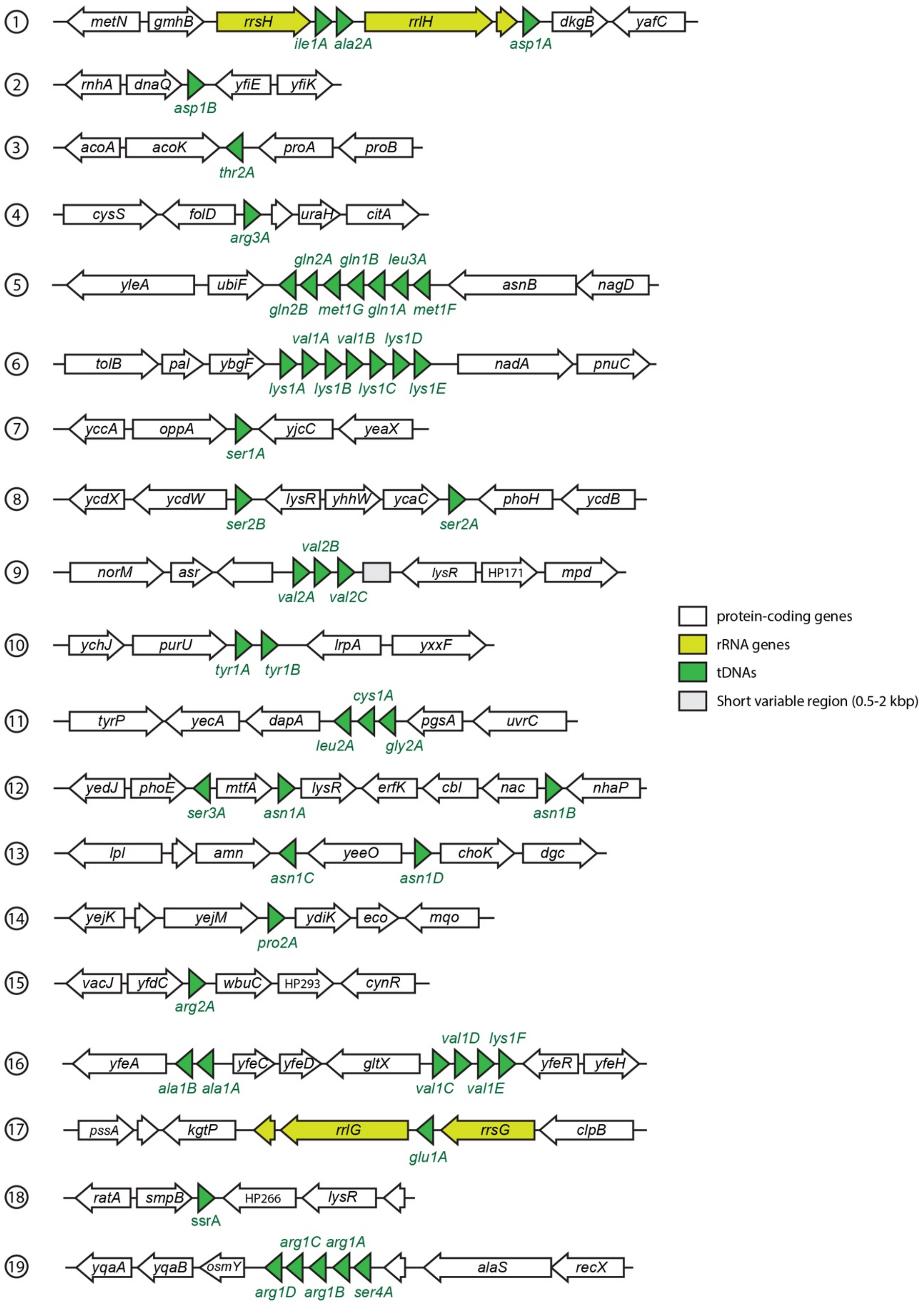
1
metN gmhB rrsH rrlH dkgB yafC
ile1A ala2A asp1A
2
rnhA dnaQ yfiE yfiK
asp1B
3
acoA acoK proA proB
thr2A
4
cysS folD uraH citA
arg3A
5
gln2A gln1B leu3A
yleA ubiF asnB nagD
gln2B met1G gln1A met1F
6
val1A val1B lys1D
tolB pal ybgF nadA pnuC
lys1A lys1B lys1C lys1E
7
yccA oppA yjcC yeaX
ser1A
8
ycdX ycdW lysR yhhW ycaC phoH ycdB
ser2B ser2A
9
val2B
norM asr lysR HP171 mpd
val2A val2C
10
ychJ purU lrpA yxxF
tyr1A tyr1B
11
cys1A
tyrP yecA dapA pgsA uvrC
leu2A gly2A
12
yedJ phoE mtfA lysR erfK cbl nac nhaP
ser3A asn1A asn1B
13
lpl amn yeeO choK dgc
asn1C asn1D
14
yejK yejM ydiK eco mqo
pro2A
15
vacJ yfdC wbuC HP293 cynR
arg2A
16
val1D lys1F
yfeA yfeC yfeD gltX yfeR yfeH
ala1B ala1A val1C val1E
17
pssA kgtP rrlG rrsG clpB
glu1A
18
ratA smpB HP266 lysR
ssrA
19
arg1C arg1A
yqaA yqaB osmY alaS recX
arg1D arg1B ser4A
protein-coding genes
rRNA genes
tDNAs
Short variable region (0.5-2 kbp)

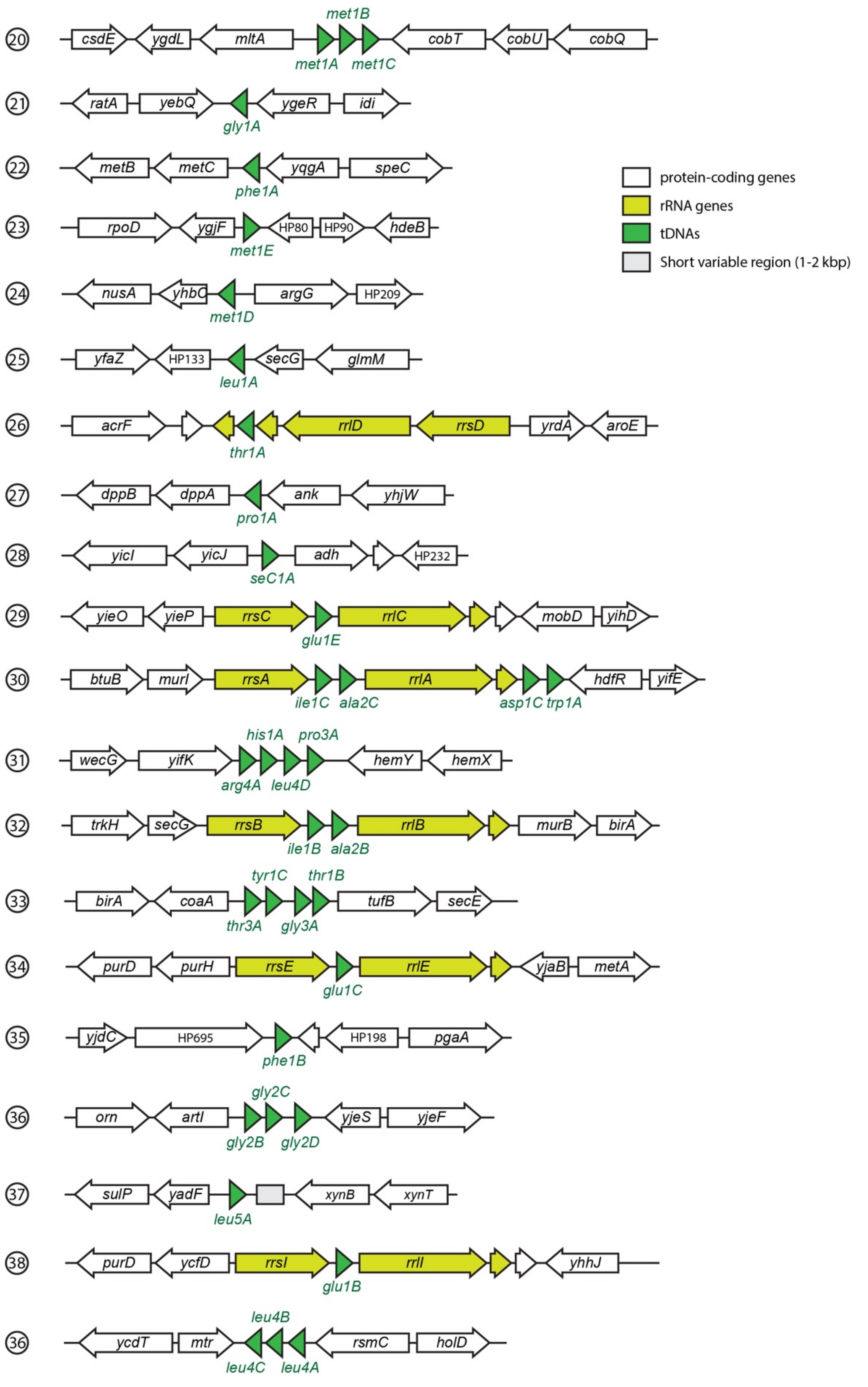


**Supplementary Figure 3. Schematic representation of the virgin contexts in which are located the different tDNAs that compose the core set of these genes in *K. pneumoniae*.**

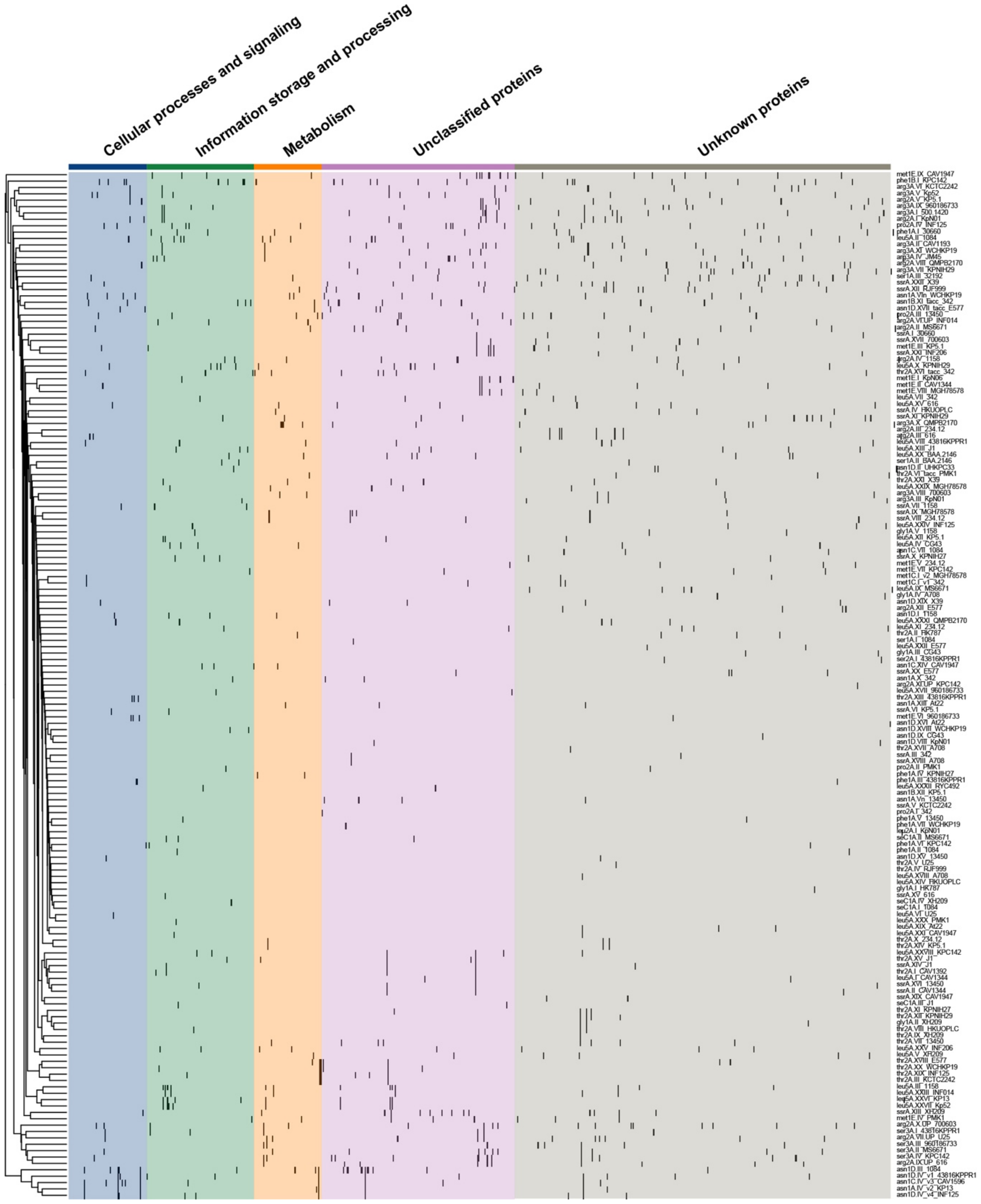


**Supplementary Figure 4. Diagram showing the presence or absence of the 3,540 GI-encoded non-redundant gene families across the different GIs identified in this study, categorized by function using the eggNOG database.** Each column represents one gene family and each row corresponds to a different GI. The clustering showed on the left was performed in base to the presence/absence of genes using the tool Roary. Black bars indicate gene presence.

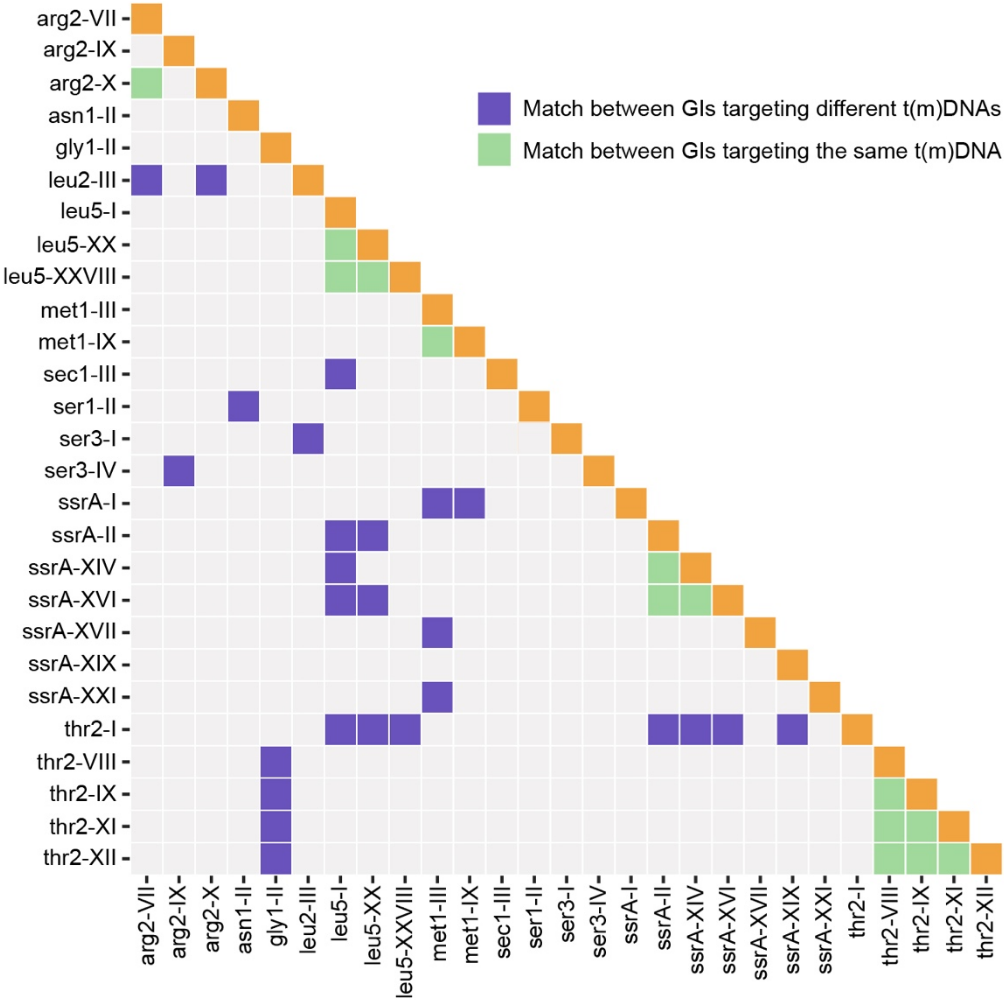


**Supplementary Figure 5. Significant matches in pairwise sequence alignments among the GIs described in this study.** The nucleotide sequence of the 162 non-redundant GIs was compared through pairwise alignments using BLASTn (a total of 13,122 different comparisons). GI pairs sharing regions showing >60% identity (e-value <0.001) are shown in green (GIs targeting the same t(m)DNA), in purple (GIs targeting a distinct t(m)DNA), or in orange (self matches).

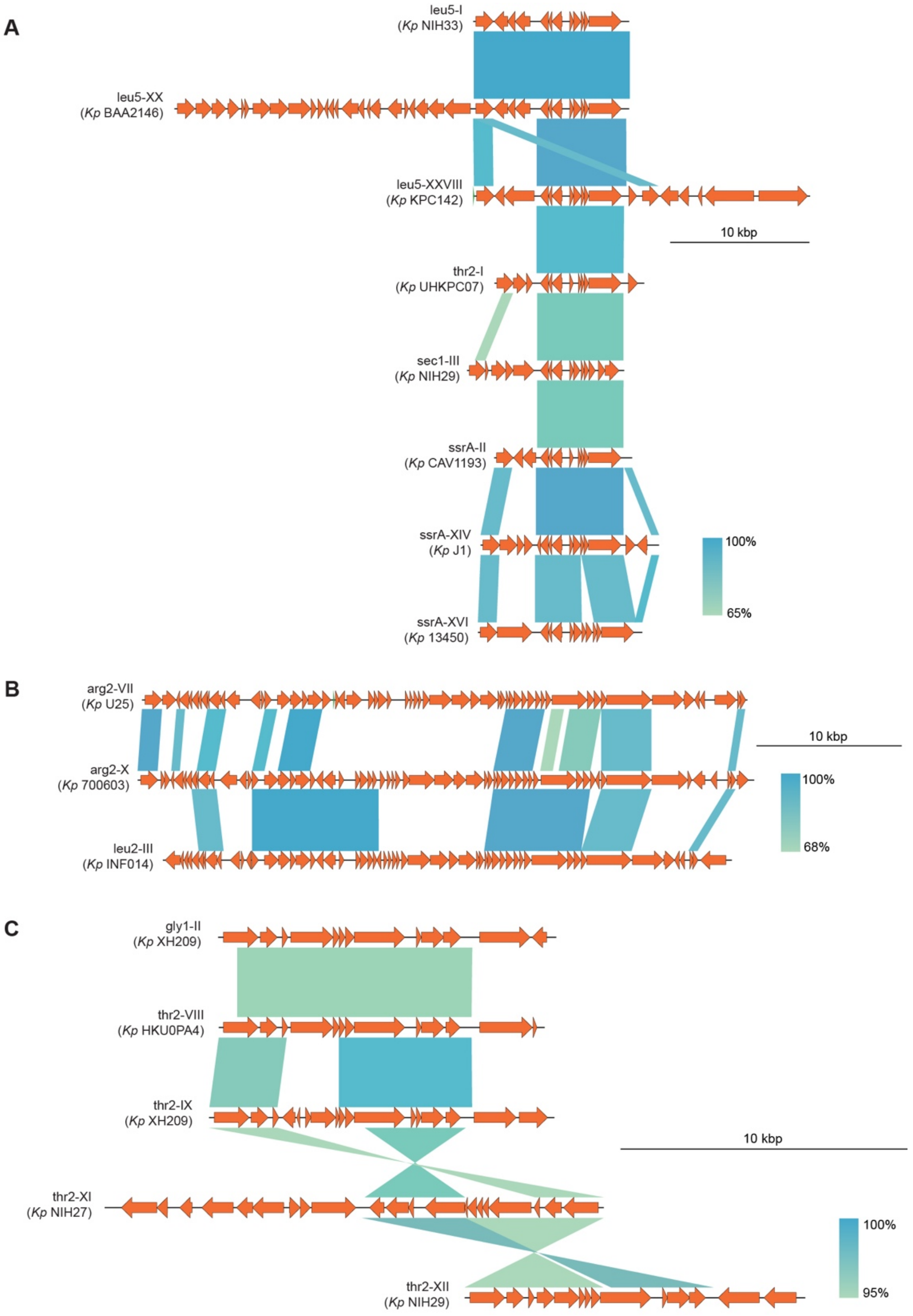
A
leu5-I
(Kp NIH33)
leu5-XX
(Kp BAA2146)
leu5-XXVIII
(Kp KPC142)
10 kbp
thr2-I
(Kp UHKPC07)
sec1-III
(Kp NIH29)
ssrA-II
(Kp CAV1193)
ssrA-XIV
(Kp J1)
100%
65%
ssrA-XVI
(Kp 13450)
B
arg2-VII
(Kp U25)
10 kbp
arg2-X
(Kp 700603)
100%
68%
leu2-III
(Kp INF014)
C
gly1-II
(Kp XH209)
thr2-VIII
(Kp HKU0PA4)
thr2-IX
(Kp XH209)
10 kbp
thr2-XI
(Kp NIH27)
100%
95%
thr2-XII
(Kp NIH29)

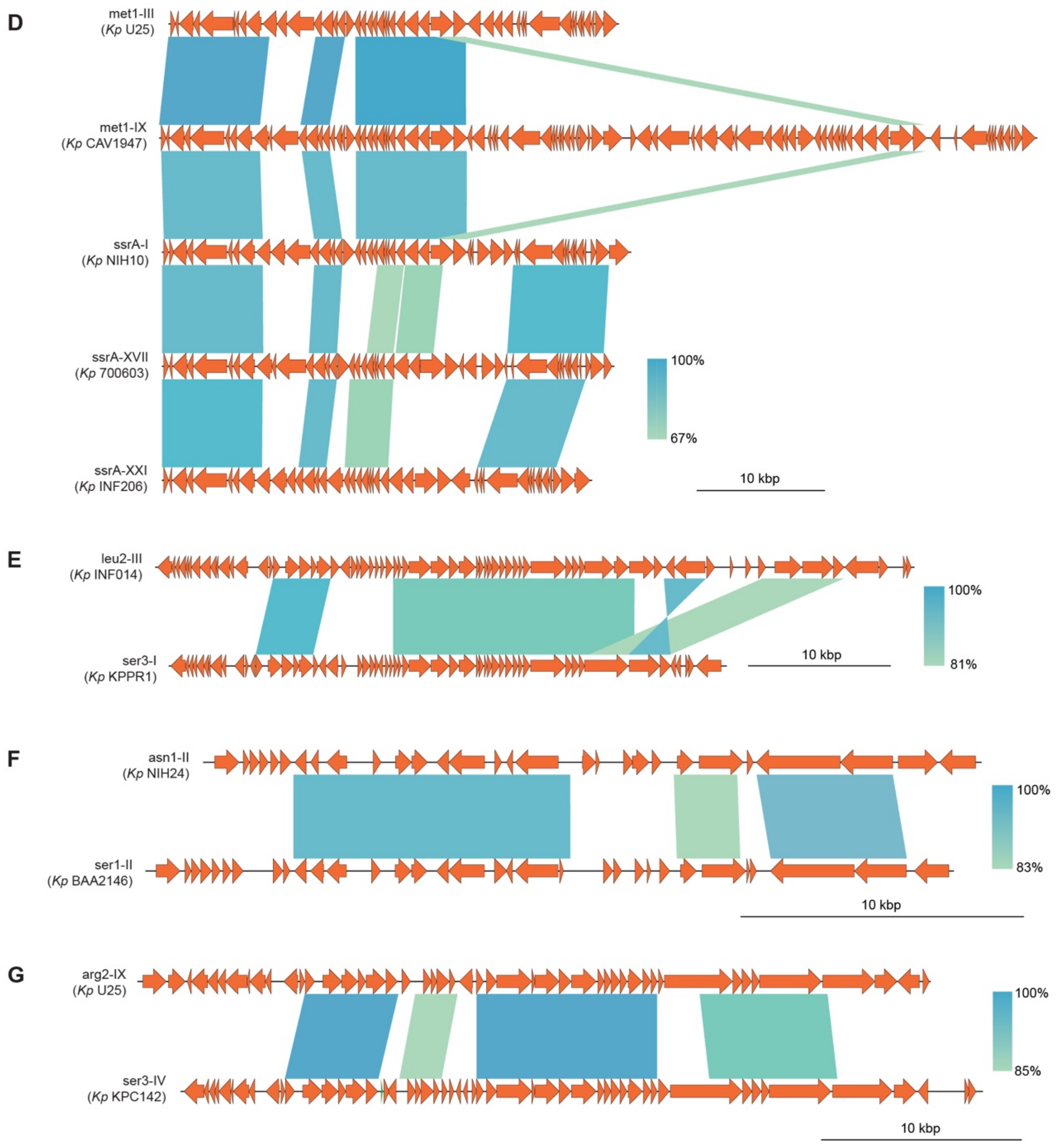


**Supplementary Figure 6. Shared DNA regions among different t(m)DNA-associated GIs found in *Klebsiella* genomes.** GIs sharing regions with >60% sequence identity are depicted including their overall gene organization. The shading in tones of green denotate the identity calculated for each shared region.

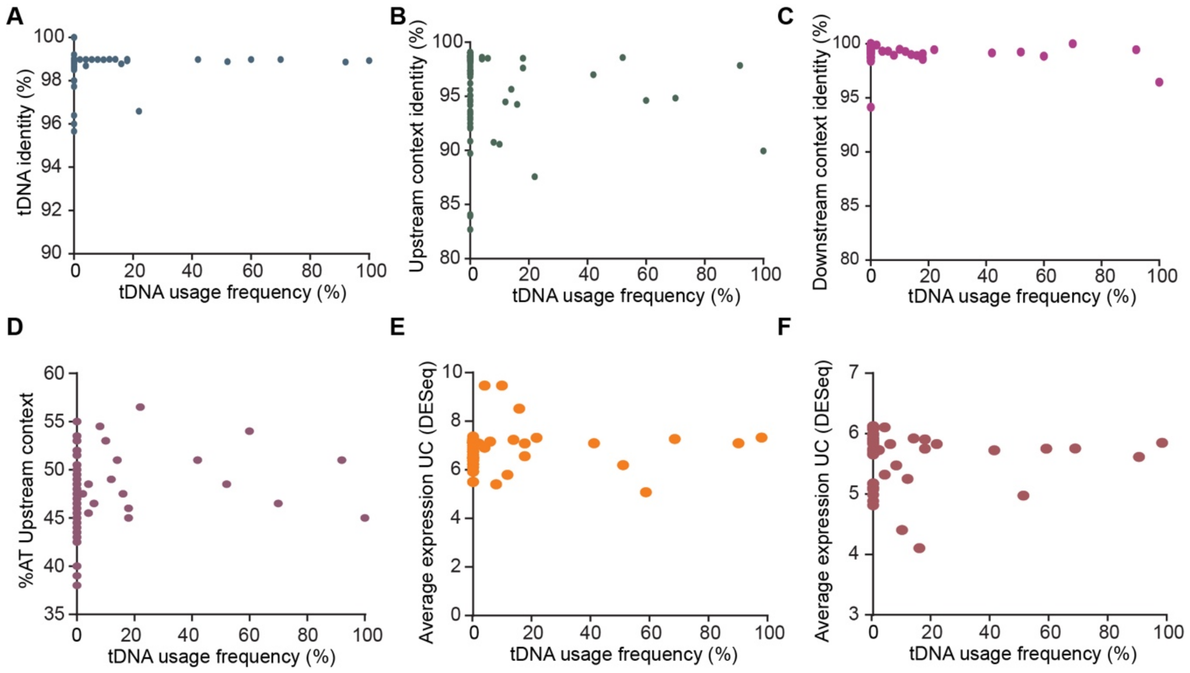


**Supplementary Figure 7.** Evaluation of the possible correlation between the usage frequency of tDNAs as integration sites and several properties of these genes and their genomic contexts. tDNA usage frequencies were compared with tDNA sequence conservation (A); the conservation of the upstream (B) and downstream (C) contexts; the AT content of the upstream context (D); and with the average expression of the genes located in the upstream (E) and downstream (F) context.

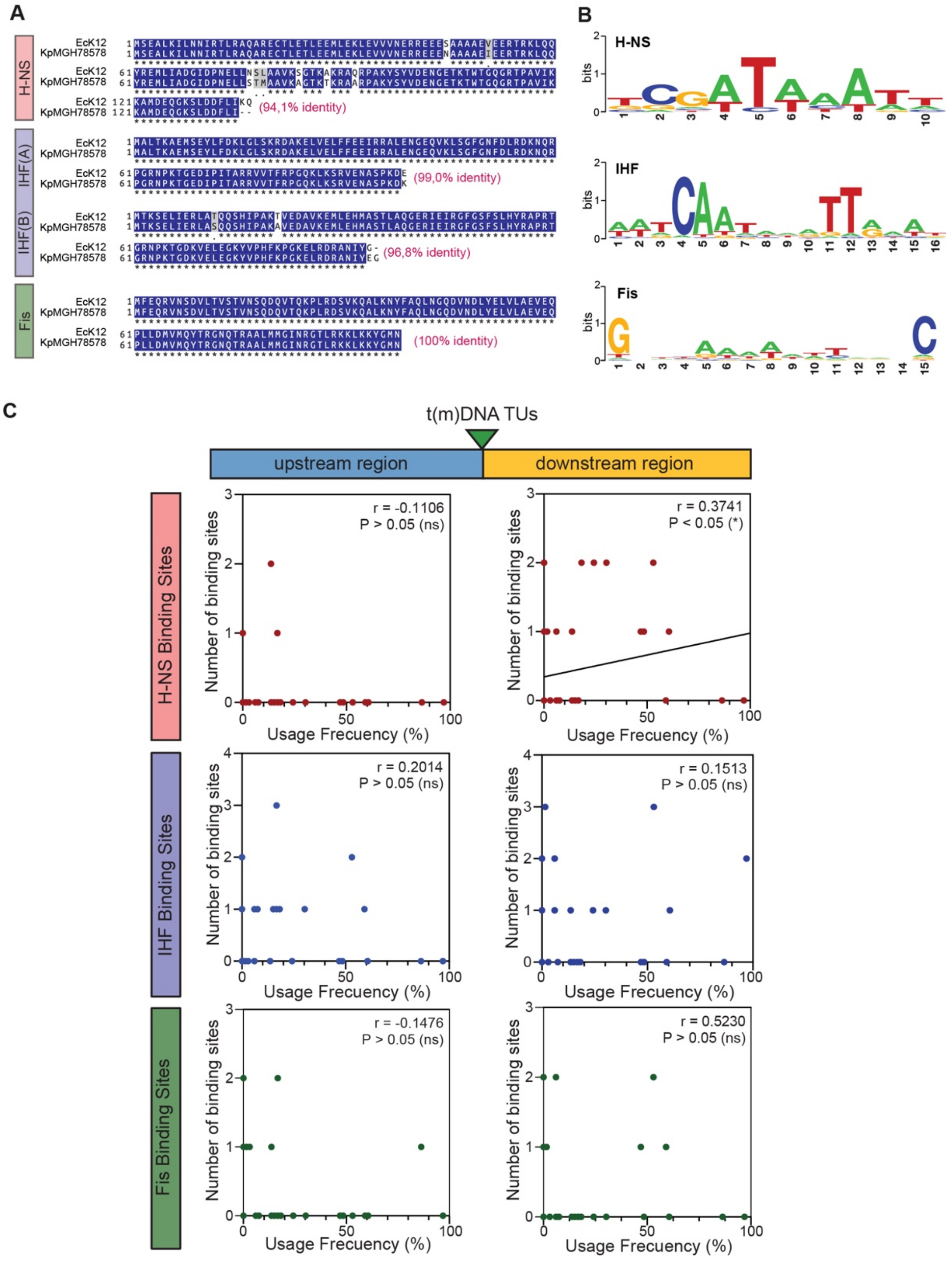


**Supplementary Figure 8.** Evaluation of the possible correlation between the usage frequency of the t(m)DNAs as integration sites and the abundance of H-NS, IHF, and Fis binding sites in the upstream and downstream regions. (A) ClustalW alignment of H-NS, IHF (subunit A and B), and Fis proteins from *K. pneumoniae* MGH78578 and *E. coli* K12. (B) LOGO plot of the consensus binding sites retrieved from the PRODORIC database and used to determine the amount of these sites present in a 4-kbp region upstream and downstream of each t(m)DNA from the *K. pneumoniae* MGH78578 chromosome. (C) Scatter plot showing for each of the 87 t(m)DNAs composing the core set, the usage frequency and the number of H-NS, IHF, or Fis predicted binding sites present in their context.

**Supplementary Table 5.** RNAseq datasets used to predict transcriptional units comprising t(m)DNAs by means of the tool rSeqTU.

| SRA ID | # of reads | Read lenght | # of bases | Reference |
|---|---|---|---|---|
| SRX119993 | 1,645,711 | 36 | 59.2M | (Kim et al., 2012) |
| SRX119994 | 757,503 | 36 | 27.3M | (9) |
| SRX5021689* | 20,023,687 | 125 | 2.5G | (10) |
| SRX5021688* | 9,937,871 | 125 | 1,2G | (10) |
| SRX5021685 | 12,142,867 | 125 | 1.5G | (10) |
| SRX5021687 | 9,446,200 | 125 | 1.2G | (10) |
| SRX5021686 | 10,783,294 | 125 | 1.3G | (10) |
| SRX5021690 | 15,394,318 | 125 | 1.9G | (10) |
| SRX5021691 | 9,043,501 | 125 | 1.1G | (10) |
| SRX5021692 | 11,258,301 | 125 | 1.4G | (10) |
| SRX814881_F | 9,247,138 | 100 | 0.9G | (11) |
| SRX814881_R | 9,247,138 | 95 | 0.9G | (11) |

*Representative datasets used for the mappings showed in Figure 3.
